# Biomolecular mechanisms for signal differentiation

**DOI:** 10.1101/2021.04.29.441952

**Authors:** Emmanouil Alexis, Carolin CM Schulte, Luca Cardelli, Antonis Papachristodoulou

## Abstract

Cells can sense temporal changes of molecular signals, allowing them to predict environmental variations and modulate their behaviour. This paper elucidates the underlying biomolecular mechanisms of time derivative computation, facilitating the design of reliable synthetic differentiator devices for a variety of applications, ultimately expanding our understanding of cell behaviour. In particular, we describe and analyse three alternative biomolecular topologies that work as signal differentiators of high accuracy to arbitrary input signals around their nominal operation. We propose strategies to preserve their performance even in the presence of high-frequency input signal components, which are detrimental to the performance of most differentiators. We found that the core of the proposed topologies appears in natural regulatory networks and we further discuss their biological relevance. The simple structure of our designs makes them promising tools for realizing derivative control action in synthetic biology.

## Introduction

Measuring the speed at which a physical process evolves over time is of central importance to science and engineering. This can be done by computing the time derivative of the function describing the process. Several examples of cellular systems exhibiting derivative action indicate that calculating the rate of change of biological processes is essential in nature. The retina of our eyes, for instance, is one of the best-studied neural networks of the brain. Its response to changes in light intensity reveals typical characteristics of derivative action which stem from the interaction between cone and horizontal cells [1, 2]. In microbiology, the chemotaxis signaling pathway in bacteria such as *Escherichia coli* involves computation of time derivatives: to navigate towards nutrients and away from toxins, bacteria are able to sample their environment as they move and convert spatial gradients into temporal ones [3–8]. Furthermore, in the context of cellular energy metabolism, *in silico* studies have revealed the role of creatine phosphate as a buffering species that allows for adaptation to a changing demand, thus exploiting the anticipatory action enabled by derivative control [9]. This observation is a specific example of a broader class of biomolecular processes where the presence of rapid buffering proves to be equivalent to negative derivative feedback [10].

In traditional engineering, differentiators refer to devices capable of applying time differentiation to an input stimulus, for example a mechanical or electrical signal. In the rapidly growing field of synthetic biology, the ability to build reliable biomolecular differentiators would offer considerable advantages [11–13]. As an immediate application, such genetic circuits would be able to track the rate of change of the concentration of biomolecules of interest, thus acting as speed biosensors. They can also allow for advanced regulation strategies in the cellular environment by enabling the construction of more efficient biocontrollers, e.g. Proportional-Integral-Derivative (PID) control schemes, the workhorses of modern technological process control applications [2]. In general, derivative control can enhance the stability of a feedback system and provide a smoother transient response.

Recent efforts in this rather underexplored research area include the design of a differentiator module consisting of linear input/output functions realized by specific processes of protein production [14, 15]. It has further been demonstrated that calculation of time derivatives is possible by using ultrasensitive topologies operating within a negative feedback loop [16], and a motif capable of computing positive and negative temporal gradients, which includes input delays and the idea of an incoherent feed-forward loop, has been presented [17]. With the aim of providing derivative action in PID control architectures, networks directly inspired by bacterial chemotaxis [18] or based on the so-called dual rail encoding have also been proposed [19, 20]. This approach avoids negativity by decomposing a signal into two nonnegative ones [21]. Finally, a derivative controller tailored to gene expression is analyzed in [22], while in the PID architecture introduced in [23], derivative control is carried out with inseparable connection to proportional and integral actions.

Here, we aim to elucidate potential mechanisms that cells exploit to achieve signal differentiation and, in parallel, to pave the way for designing efficient and reliable artificial signal differentiator devices in a cellular context. Notably, we address commonly encountered issues related to guaranteeing satisfactory accuracy of temporal derivative calculation for arbitrary molecular signals. We also focus on motifs that can function as independent, general-purpose differentiators without being limited to specific roles such as control strategies. Moreover, the theoretical assumptions under which these motifs are able to work as intended can be practically satisfied in designs.

Specifically, we introduce three biomolecular architectures capable of functioning as signal differentiators of high accuracy around their equilibria. We call them <u>Bio</u>molecular <u>S</u>ignal <u>D</u>ifferentiators (BioSD). Each of these networks can be interpreted as a modular and tunable topology inside the cell that accepts a molecular signal as an input and produces an output signal proportional, or ideally equal, to the time derivative of the input signal (Fig. 1a). The output corresponds to a biochemical species, whose concentration can be measured. The proposed architectures provide simple blueprints for the design of synthetic biomolecular differentiators, but can also be interpreted as lenses through which derivative action in natural systems can be identified and studied.

**Fig. 1:**
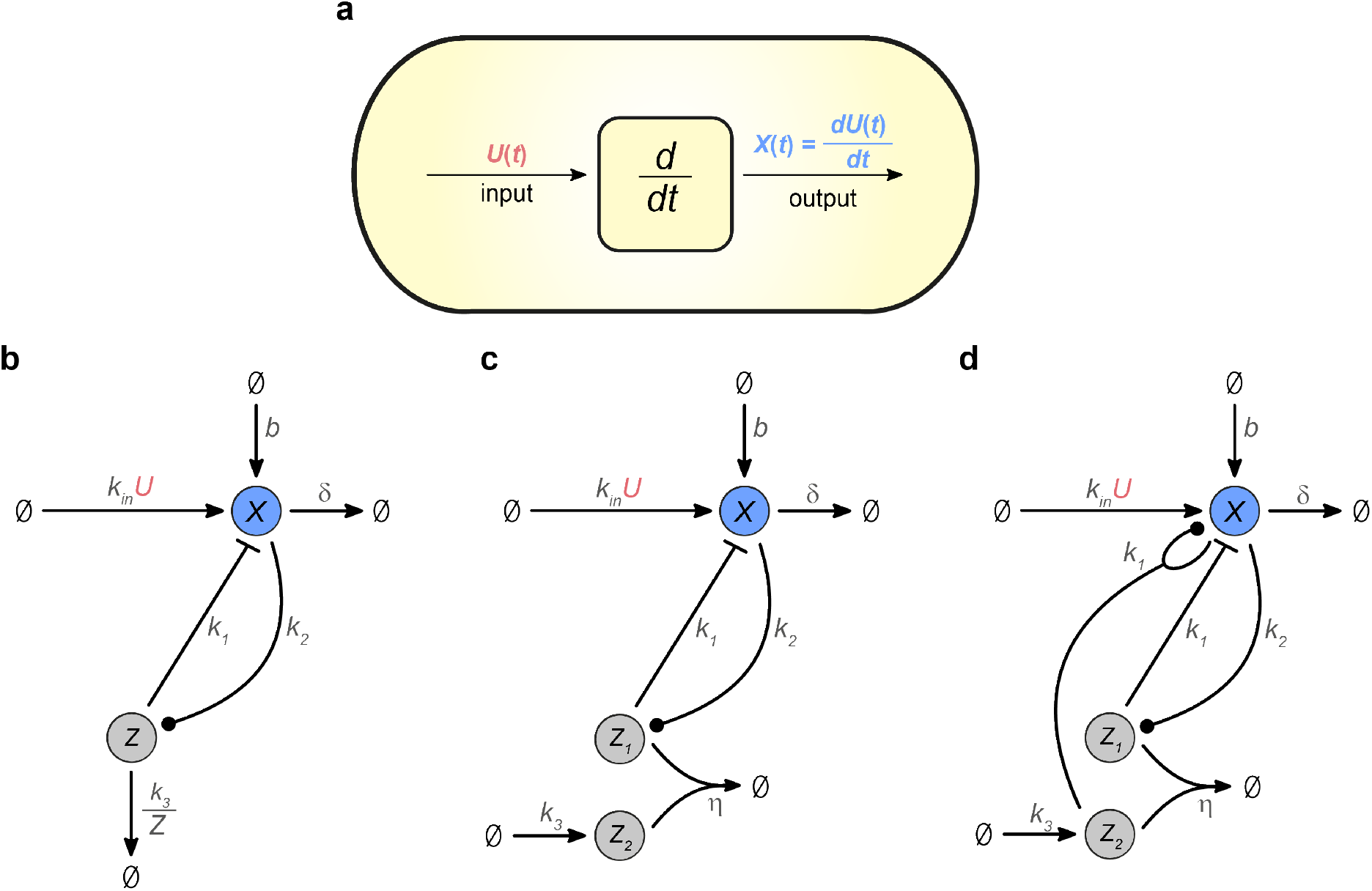
Biomolecular structures capable of signal differentiation. Schematic representation of **a** the notion of signal differentiation carried out by a biomolecular device inside the cell, **b** Biomolecular Signal Differentiator −1 (BioSD-I), **c** Biomolecular Signal Differentiator - II (BioSD-II), and **d** Biomolecular Signal Differentiator - III (BioSD-III). In **b**, **c**, **d** the following notation is adopted: (→) means that the transformation of reactants into products only happens in the direction of the arrow. 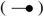 indicates that reactants enable product formation without being consumed. 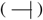 denotes inhibition of products by a reactant where the reactant is not consumed.

We demonstrate the special characteristics of the three BioSD architectures (BioSD-I, II, and III) via theoretical analyses and numerical simulations. We also discuss the major limitation of both technological and biological differentiators, namely amplification of undesired high-frequency components of the input signal, and propose strategies to overcome this obstacle. Finally, we show the occurrence of one of the BioSD topologies in natural regulatory networks involved in bacterial adaptation to stress conditions, highlighting the biological relevance of the presented designs.

## Results

### Biological structure

We begin by presenting the molecular interactions in the BioSD circuits as chemical reaction networks (CRNs). These circuits represent three alternative topological entities which, under certain assumptions, realize the same concept of signal differentiation. In the analysis that follows, the input and output signal of the differentiators are modeled as biomolecular species, namely *U* and *X* respectively.

Figure 1b illustrates the first architecture, BioSD-I, which consists of the following reactions:

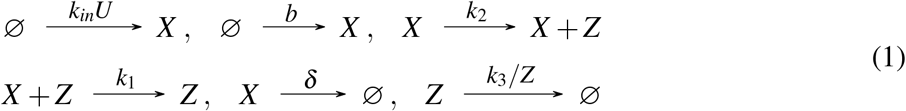

Here, the production of output species *X* depends on two reactions. One of them has a constant rate while the other occurs at a rate proportional to the concentration of input species *U. X* also catalyzes the formation of species *Z* which, in turn, inhibits *X*. Finally, the removal rate of *X* is proportional to its concentration while *Z* adheres to a constant rate of decay. One way to attain this behaviour is through enzyme-catalyzed degradation of *Z* where the enzyme is operating at saturating substrate levels (see Supplementary Note 1.1).

In the second architecture, BioSD-II (Fig. 1c), the formation process of output species *X* is the same as in BioSD-I while *Z*_1_, the production of which is facilitated by *X*, and *Z*_2_ annihilate each other. *Z*_1_ inhibits *X* which decays in the same way as in BioSD-I. The reactions that form the corresponding CRN are:

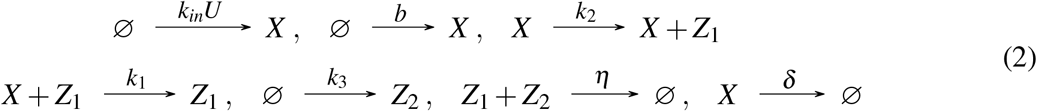

Finally, Fig. 1d shows the third topology, BioSD-III, which is described by the reactions:

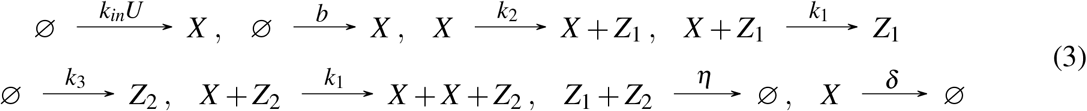

This CRN includes an autocatalytic-like reaction: *X* is able to produce more of itself in the presence of *Z*_2_. The rest of its structure is identical to the CRN of BioSD-II.

### Mathematical description

We now derive the dynamics of the proposed BioSD networks using the law of mass action [24] unless otherwise stated, adopting the same order of presentation as in the preceding section.

BioSD-I (CRN (1)) can be described by the following system of ordinary differential equations (ODEs):

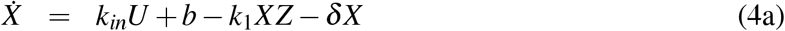

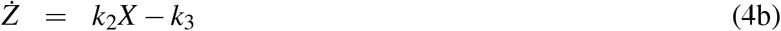

Note that the enzymatic degradation of *Z* is assumed to follow Michaelis-Menten kinetics, as previously discussed.

Next, from CRN (2) we obtain the following ODE model for BioSD-II:

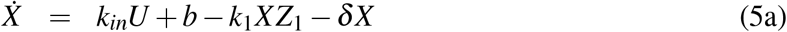

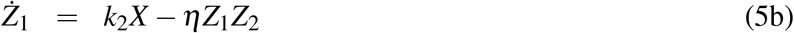

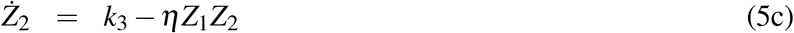

For the last circuit, BioSD-III, CRN (3) can be modelled using the following ODEs:

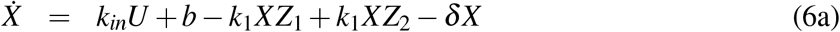

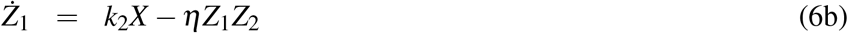

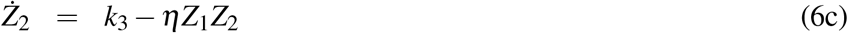

We can prove that for a constant input *U*\*, each of the BioSD network models has a unique locally exponentially stable equilibrium (see Supplementary Note 1). Near their steady-states, the circuits are able to differentiate biological signals with high accuracy, as shown in the next section. Finally, for the purpose of this study we assume that the parameter *η* in BioSD-II is sufficiently large (a notion mathematically defined in Supplementary Note 2). This implies that the binding between species *Z*_1_ and *Z*_2_ occurs with sufficiently high affinity. This constraint does not have to hold for BioSD-III, which includes the same annihilation reaction.

### Achieving biological signal differentiation

In order for the proposed biomolecular modules to work as signal differentiators, we desire for their output *X* to be equal or at least proportional to the derivative of their input *U*. This immediately raises the following challenge: both *U* and *X* refer to biomolecular species concentrations and, by extension, represent non-negative signals. However, in the general case, the derivative of a nonnegative signal can take negative values and, as a result, *X* would need to go below zero. Thus, it could be argued that *X* is unable to express the rate of change of an arbitrary input signal. An obvious way to overcome this obstacle is to add a bias to the computed derivative. As we demonstrate here, the perfect candidate for realizing this bias is the steady state of *X* around which derivative action can be achieved. A detailed mathematical derivation of the results that follow can be found in Supplementary Note 3.

We are interested in the local behaviour of the BioSD networks and therefore consider input stimuli that do not force them to operate far away from their equilibrium. Subsequently, we assume that every input signal can be described as:

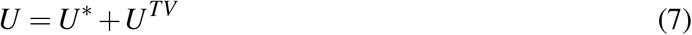

where *U*\* is constant while *U^TV^* is time-varying.

We now establish conditions for accurate signal differentiation. For this purpose, we introduce a non-dimensional parameter:

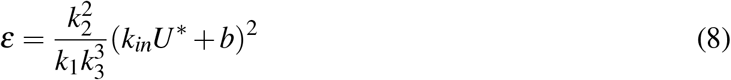

It can be shown that if *ε* is close to zero, i.e.:

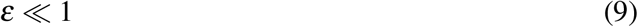

then the local dynamics of *X* in BioSD-I (Eqs. (4a)–(4b)), BioSD-II (5a)–(5c)) and BioSD-III (Eqs. (6a)–(6c)) can be approximated by:

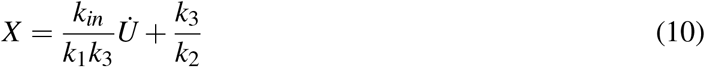

In fact, there is a family of input signals for which the BioSD topologies are able to provide accurate differentiation without the need of satisfying condition (9). More specifically, for input signals for which the term *U^TV^* in Eq. (7) is of the form:

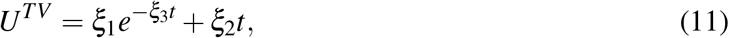

where *ξ*_1_, *ξ*_2_ are arbitrary constants and 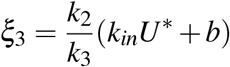, we can derive Eq. (10) for all three networks without selecting an appropriate parameter combination to drive *ε* close to zero.

From Eq. (10), we can see that the BioSD modules use the biomolecular concentration 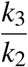 as a bias. Around this point they can operate as signal differentiators, producing an output signal component which is proportional to the derivative of the input or even equal if we can ensure *k_in_* = *k*_1_*k*_3_. The bias therefore depends only on two parameters which, ideally, can be adjusted as desired. This provides us with the freedom of choosing any (fixed) concentration of *X* as a bias, which will remain unchanged regardless of the rest of the model parameters, the input stimulus, or potential constant disturbances on the output. To appreciate this further, we recall the birth reaction for *X* with constant rate *b*, which is included in each of the proposed CRNs. Besides its role as a structural requirement, this birth reaction can also represent an external constant disturbance applied on *X*; this, however, does not affect the zero-level we choose for our measurements. Once the concentration of *X* reaches this level, it will stay there until an input excitation appears and it will come back once the excitation stops. Hence, the previously mentioned fixed concentration can also be seen as a “rest position” for the differentiators.

The feature just described is of key importance and stems mainly from the following two sources: the (input-to-state) stability that characterizes BioSDs and the fact that the steady-state of the output coincides with the aforementioned zero-level concentration. The latter is achieved due to integration carried out by the a ‘memory’ function which is realized via species *Z* within BioSD-I and the (not physical) quantity *Z*_1_ – *Z*_2_ within BioSD-II, III.

### Tunability and accuracy

It is convenient for the circuit designer who aims to implement the BioSD topologies to be able to choose the parameter values and ensure that the resulting differentiators meet the expected performance requirements. Nonetheless, there may be cases where the number of system parameters that can be suitably tuned is limited, for instance due to constraints related to the cellular processes involved in the circuits under investigation. Even in this case the architecture of our circuits allows for some tunability as long as the designer can choose some crucial parameters.

Consider for example the extreme scenario where only one of the model parameters can be regulated. If this parameter is *k*_3_, then, according to Eq. (10), its appropriate tuning may result in an acceptable gain by which the output signal is multiplied (output gain) and bias based on which this signal is measured. At the same time, Eq. (8) reveals that a small change in *k*_3_ can affect *ε* significantly and thereby make it sufficiently small as Eq. (9) commands.

It immediately emerges from the above that the way we tune the BioSD networks defines the level of accuracy regarding their derivative action. Indeed, *ε* is subject to almost all parameter rates in these networks and, as pointed out in the previous section, an *ε* close to zero is a fundamental requirement if our goal is to construct differentiators that work accurately for all kinds of input stimuli. In fact, in Supplementary Note 4 we show for arbitrary input signals that the ability of the output to track the ideal derivative of the input improves as *ε* approaches zero.

### Sensing the response speed of biomolecular networks

We now demonstrate through an example the capacity of BioSD modules to compute the temporal derivative of biological signals. At the same time, we highlight one of their potential applications discussed above, namely as rate-of-change detectors or speed biosensors.

We consider the antithetic motif [18, 25–30]:

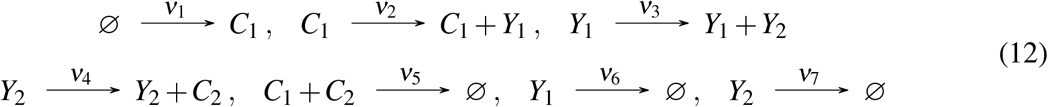

Species *Y*_1_, *Y*_2_ represent an arbitrary biological process whose output, *Y*_2_, can be robustly steered towards a desired value 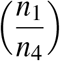. This is feasible through the feedback integral control which is implemented via species *C*_1_, *C*_2_, thus achieving robust perfect adaptation. Depending on the parameter rates, the dynamics of the above architecture can be either stable or unstable. Nonetheless, even in a stable system, the species of interest, *Y*_2_, sometimes displays a long-lasting transient response with damped oscillations before it settles to a steady-state. This provides an opportunity to assess the ability of the BioSD networks to calculate the speed at which these oscillations evolve.

In order for a BioSD device to function as a biosensor for CRN (12), a suitable interconnection between these circuits is required while preserving the modularity [24] of the two networks and avoiding any loading problems. One way to accomplish this is through the reaction:

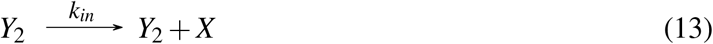

where *Y*_2_ plays the role of the input species *U* without being consumed. Alternatively, in case the nature of *Y*_2_ prevents it from directly producing *X*, we can use a separate sensory species *S* which is capable of participating in the formation of *X* and it is co-expressed with *Y*_2_, i.e.:

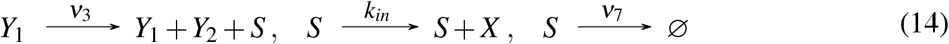

Adopting the second interconnection as the most general one, we manifest in Fig. 2 that the rate of change of the concentration of *Y*_2_ is accurately represented by the output of the BioSD networks provided that *ε* is sufficiently small (condition (9)). As already discussed, satisfying this constraint is necessary unless the input of the differentiators belongs to the family of signals defined by Eqs. (7), (11), which is clearly not true in this case.

**Fig. 2:**
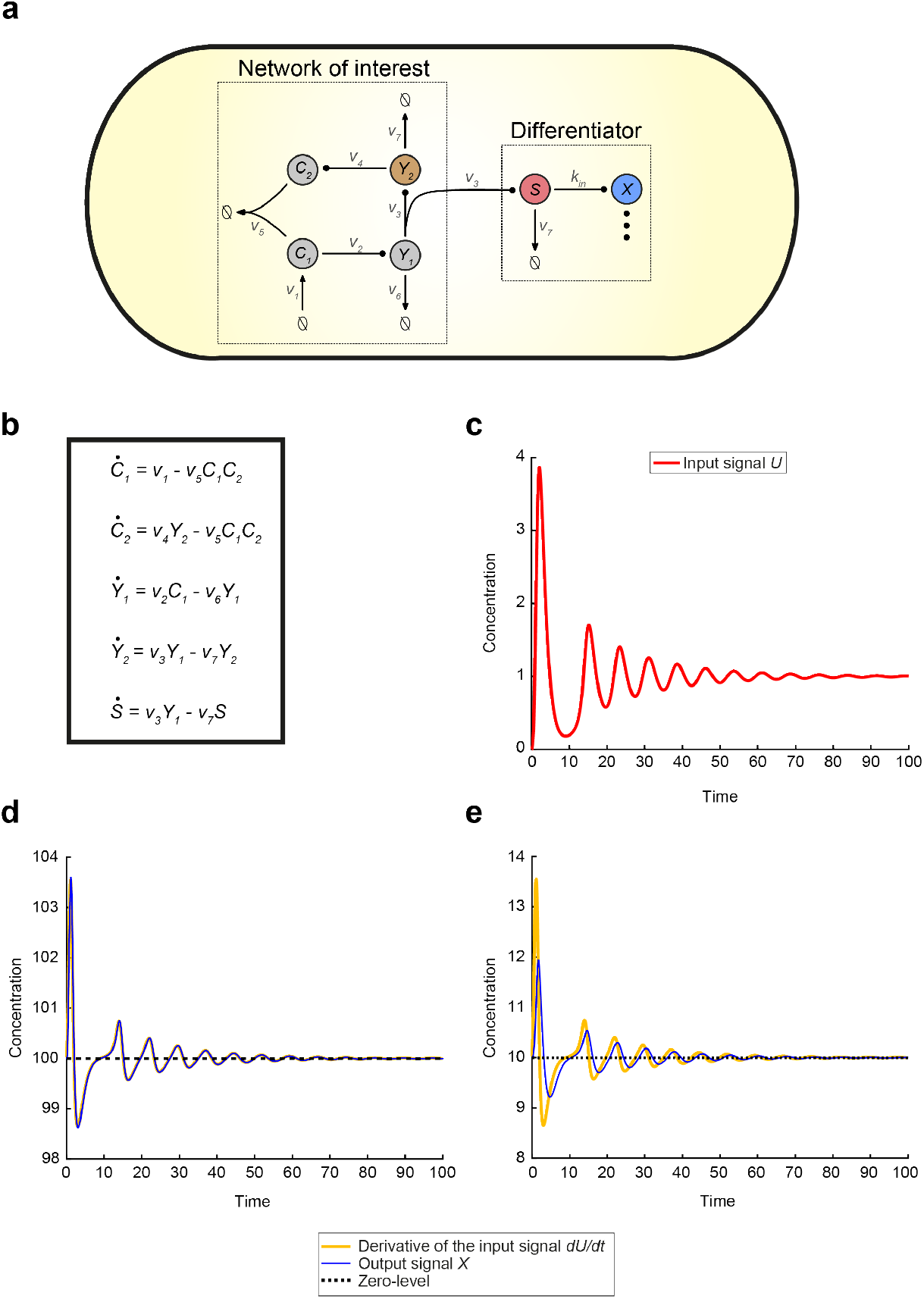
Sensing the rate-of-change of a synthetic regulatory biomolecular network through a Biomolecular Signal Differentiator. **a** Schematic of CRN (12) (network of interest) accompanied by a BioSD device (differentiator) which measures the speed of the output, *Y*_2_ of the network via the sensing mechanism in (14). We adopt the same arrow notation as in Fig. 1 while the symbol 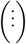 represents any of the three BioSD devices. **b** ODE model capturing the dynamics of the topology given by (12), (14). As anticipated, the behaviour of species *Y*_2_ and *S* is described by the same equation. **c** Input *U* of the differentiator coincides with species *S* and results from the simulation of the ODE model depicted in **b** with the following parameter values: *ν*_1_ = *ν*_2_ = *ν*_4_ = 2, *ν*_3_ = 4, *ν*_5_ = 12, *ν*_6_ = *ν*_7_ = 1. **d** Simulation of BioSD-I (Eqs. (4a), (4b)) response to the input shown in **c** using the following parameter values which satisfy condition (9): *k_in_* = *k*_3_ = *b* = 100, *k*_1_ = *k*_2_ = 1, *δ* = 0.5. As can be seen, the output, *X*, of the differentiator is an accurate replica of the derivative of input *U*. **e** The simulation in **d** is repeated after replacing the value of both *k_in_* and *k*_3_ with 10. This change leads to violation of condition (9). In fact, *ε* ≫ 1 and, thus, poor performance of the differentiator is observed. The respective simulations regarding BioSD-II and BioSD-III are presented in Supplementary Note 7. As expected, their responses are identical to those of BioSD-I.

For comparison purposes, we now replace the circuit (12) with the general birth-death process:

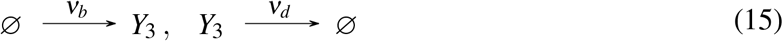

maintaining the same kind of interconnection, as illustrated in Fig. 3. Focusing on the linear regime of its response (which is obviously aligned with Eq. (11)), it can be seen that, although *ε* is much larger than unity, BioSD networks are now able to provide accurate signal differentiation.

**Fig. 3:**
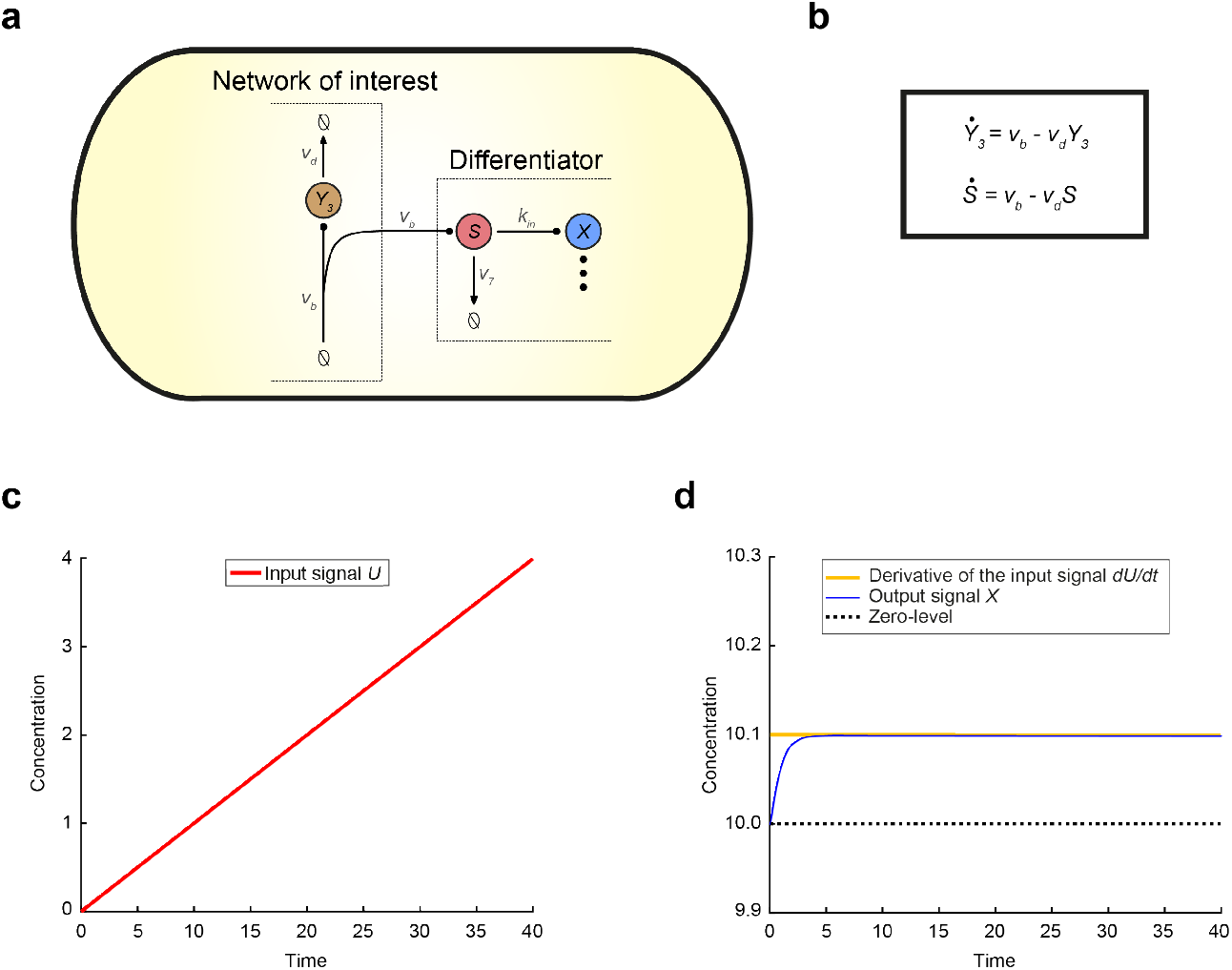
Sensing the rate-of-change of a birth - death biomolecular process through a Biomolecular Signal Differen-tiator. **a** Schematic of CRN (15) (network of interest) accompanied by a BioSD device (differentiator), which measures the speed of the output of the network (*Y*_3_) via the sensing mechanism in (14). We adopt the same arrow notation as in Fig. 1 while the symbol 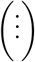 represents any of the three BioSD devices. **b** ODE model capturing the dynamics of the topology given by (15), (14). As anticipated, the behaviour of species *Y*_3_ and *S* is described by the same equation. **c** Input *U* of the differentiator coincides with species *S* and results from the simulation of the ODE model depicted in **b** with the following parameter values: *ν_b_* = 0.1, *ν_d_* = 0.001. **d** Simulation of the BioSD-I (Eqs. (4a), (4b)) response to the input presented in **c** using the following parameter values: *k_in_* = *k*_3_ = 10, *b* = 100, *k*_1_ = *k*_2_ = 1, *δ* = 0.5 (same as in Fig. 2e). Although *ε* ≫ 1 (condition (9) is violated), the output, *X*, of the differentiator is now an accurate replica of the derivative of input *U*. This is due to the fact that the input *U* shown in **c** belongs to the class of signals defined by Eqs. (7, 11). The respective simulations regarding BioSD-II and BioSD-III are presented in Supplementary Note 7. As expected, their responses are identical to those of BioSD-I.

### Response to input signals corrupted by high-frequency noise

Potentially the most important problem of differentiator devices is their sensitivity to high-frequency noise components which the applied input signal may contain [2]. To this end, we consider an input signal with a time-varying component^1^

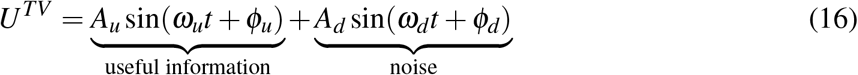

where the actual signal we want to differentiate (useful information) is accompanied by undesired fluctuations (noise) arising, for instance, from unintended cross-talk interactions [24]. Assuming perfect differentiation, we get:

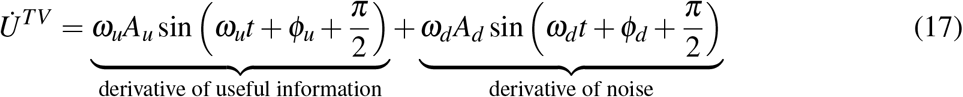

Hence, even if the level of input corruption is low (e.g. *A_d_* is much smaller than *A_u_* - Eq. (16)), the damage in the output of a perfect differentiator may be detrimental in case of a rapidly fluctuating noise signal (*ω_d_* high). That is, *ω_d_A_d_* can be made arbitrarily large compared to *ω_u_A_u_* (Eq. (17)) and, therefore, it is possible for the derivative of the useful signal to be completely drowned out by the derivative of some high frequency input noise.

Interestingly, the BioSD topologies allow us to deal with this noise amplification by suitably adjusting *ε* (see Supplementary Note 5 for more details). It has been emphasized already that the accuracy of derivative action drops as this parameter moves away from zero. However, the intensity of this phenomenon can vary significantly depending on the frequency content of each input signal. In fact, for a given nonzero *ε*, there is a range of frequencies where signal differentiation can be successfully performed while over a different range of higher frequencies signal attenuation is carried out instead (Fig. 4a). Nevertheless, between these regions of the frequency spectrum, neither differentiation nor attenuation (at least of satisfactory accuracy) may be achieved (Fig. 4b).

**Fig. 4:**
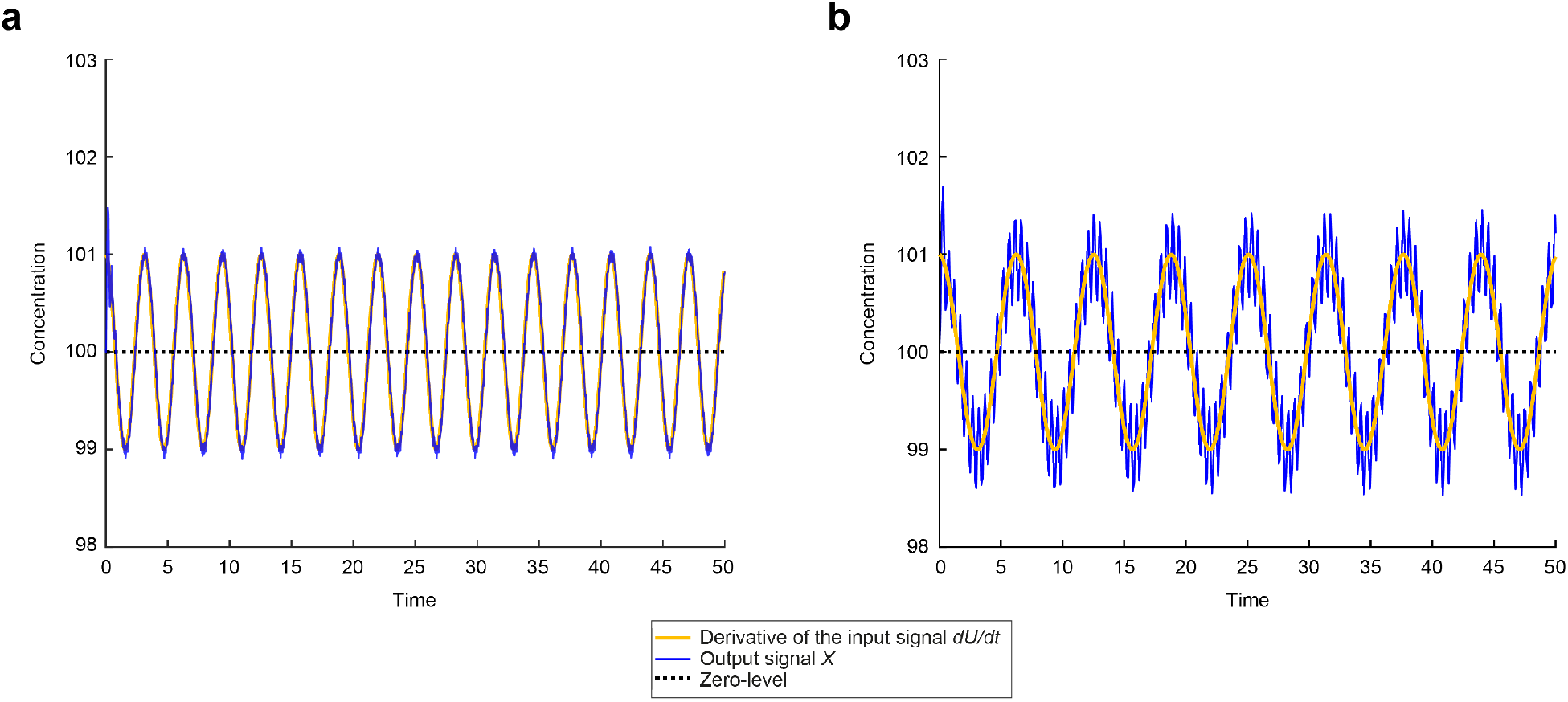
Response of Biomolecular Signal Differentiators to input signals with undesired high frequency components. Without loss of generality we select BioSD-I (Eqs. (4a), (4b)) to plot: **a** Simulated response to an input of the form (7), (16) using the following parameter values: *U*\* = 1.2, *A_u_* = *ω_u_* = 1, *A_d_* = 0.2, *ω_d_* = 400, *ϕ_u_* = *ϕ_d_* = 0, *k_in_* = *k*_3_ = *b* = 100, *k*_1_ = *k*_2_ = 1, *δ* = 0.5 (condition (9) is satisfied). Consequently, with respect to the input signal, the frequency of the undesired component (noise) is 400 times higher than that of the component of interest (useful information). It is evident that significant noise attenuation takes place and the accuracy of signal differentiation therefore remains very high. **b** The simulation in **a** is repeated after changing the value of *ω_d_* to 50 which makes the noise 40 times faster compared to the useful information. As can be seen, there is a decrease in the accuracy level of signal differentiation since the input noise of this frequency cannot be filtered adequately.

### A structural addition for enhanced performance

In case there are increased requirements for noise reduction that cannot be easily met via parameter tuning, we present an alternative version of the BioSD networks with higher noise insensitivity, which we call BioSD^*F*^ (Fig. 5a). These topologies are described by the same CRNs presented in the section **Biological structure**, but amended appropriately.

**Fig. 5:**
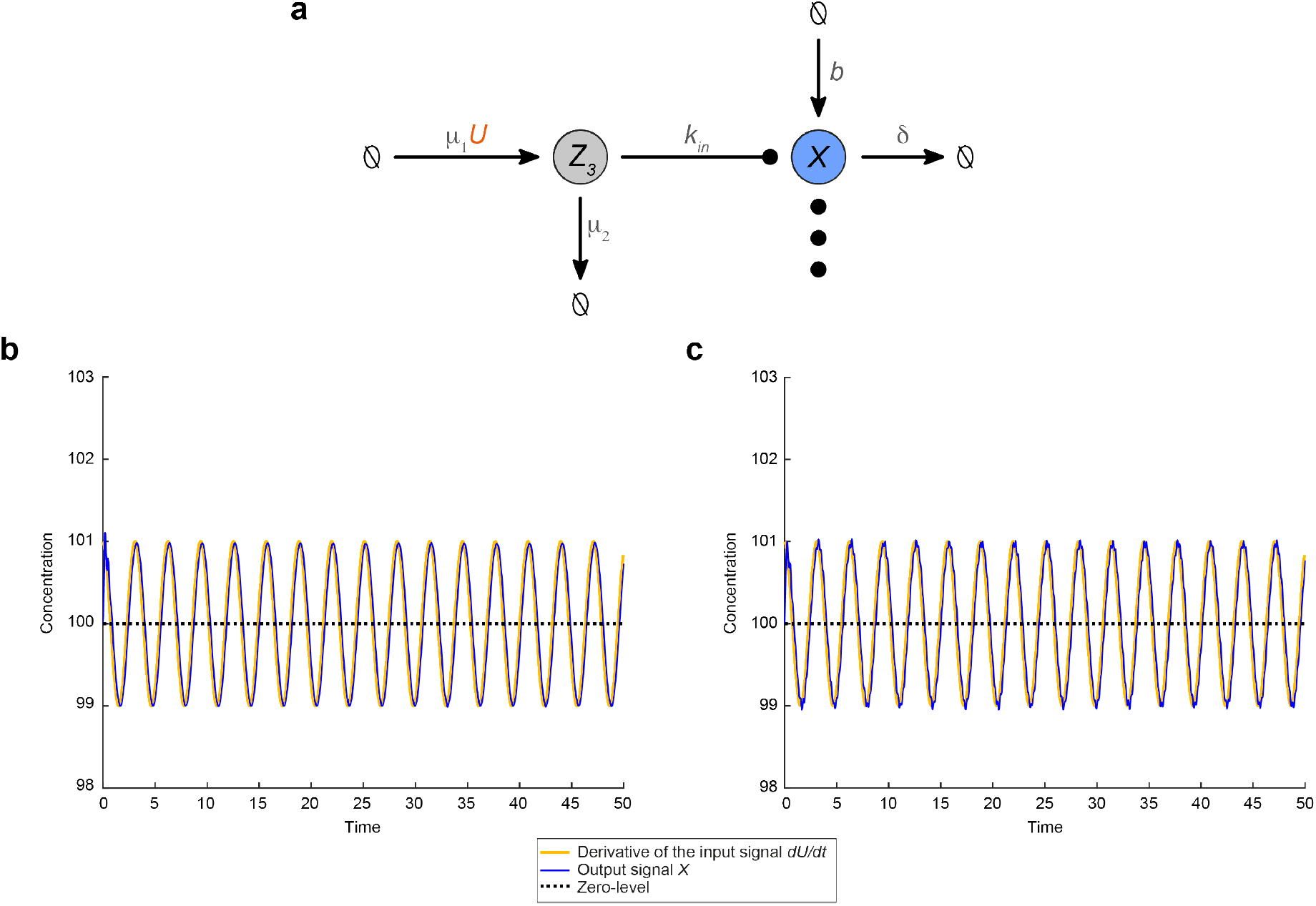
An alternative version of Biomolecular Signal Differentiators with an enhanced capability of input noise filtering. **a** Schematic structure of BioSD^*F*^. We adopt the same arrow notation as in Fig. 1 while the symbol 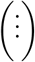 represents the remaining reactions composing any of the three BioSD devices. For comparison purposes, we focus on a BioSD^*F*^ device based on BioSD-I to re-plot the simulation of **a** Fig. 4a and **b** Fig. 4b for the same values of the mutual parameters and *μ*_1_ = *μ*_2_ = 5. It is apparent that in both **a**, **b** very strong input noise attenuation takes place and the differentiation of the useful signal is thus conducted with significantly high accuracy.

Recalling CRNs (1), (2), (3), we see that input signals are applied to BioSD modules through the reaction:

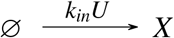

In BioSD^*F*^ topologies, the above is replaced by the following set of reactions:

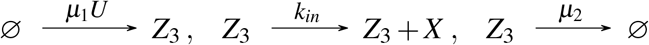

The additional species *Z*_3_ is produced by the input species and degrades in the traditional manner while it catalyzes the formation of the output species. This structural addition is inspired by the work in [31, 32], where biomolecular concepts from the area of signal processing were studied. In the following, we briefly present the main features of BioSD^*F*^ modules – a comprehensive analysis of their behaviour can be found in Supplementary Note 6.

There is a band of frequencies where, under the assumptions discussed in **Achieving biological signal differentiation locally**, the output of BioSD^*F*^ networks can be approximated by:

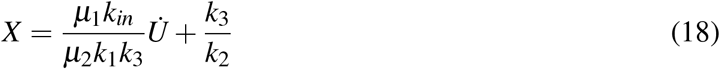

Compared to the original BioSD topologies (Eq. (10)), we now have two additional tuning parameters (*μ*_1_, *μ*_2_) with respect to the output gain. However, the major advantage of this version of differentiators is an enhanced capability of noise filtering. In fact, we can have a greatly extended frequency range across which very strong attenuation of high frequency input noise can be achieved (Figs. 5b, 5c). At the same time, the width of this frequency band depends on *μ*_2_ and can be adjusted appropriately.

### Biomolecular Signal Differentiators in natural regulatory networks

As outlined in the introduction, derivative action appears to be an important mechanism in various biological systems. To explore the biological relevance of the proposed BioSDs for cellular adaptations to environmental changes, we identified two naturally occurring regulatory network motifs resembling the BioSD-II network. Note that these natural topologies are operating in the larger context of complex regulatory networks involving a plethora of signaling factors, some of which remain to be identified. We therefore describe the relevant motifs but do not comprehensively detail all interactions occurring in the biological system. In addition to the natural regulatory networks described here, we outline possible synthetic implementations for all BioSD networks in Supplementary Note 8.

#### Stationary phase and starvation response – RpoS regulatory network

As shown in Fig. 6a, we found the BioSD-II motif in the context of adaptation to nutrient starvation and entry into stationary phase, which is mediated by the sigma factor RpoS in *E. coli* and related bacteria (reviewed in [33, 34]). Stress conditions, such as nutrient depletion or high pH, serve as the input *U*. While RpoS is present at low levels (*b*) in exponentially growing cells, its expression is significantly increased through both transcriptional and post-transcriptional regulation in response to environmental stresses or starvation [33]. One of the genes whose expression is dependent on RpoS is *rssB*, which encodes a response regulator. RssB binds to RpoS and mediates its degradation by the ClpXP protease [35], thus functioning as *Z*_1_. Nutrient starvation also induces the expression of several anti-adaptor proteins (Ira; <u>i</u>nhibitor of <u>R</u>ssB <u>a</u>ctivity). These proteins bind to RssB and prevent RpoS degradation [36], which corresponds to the action of *Z*_2_ in BioSD-II.

**Fig. 6:**
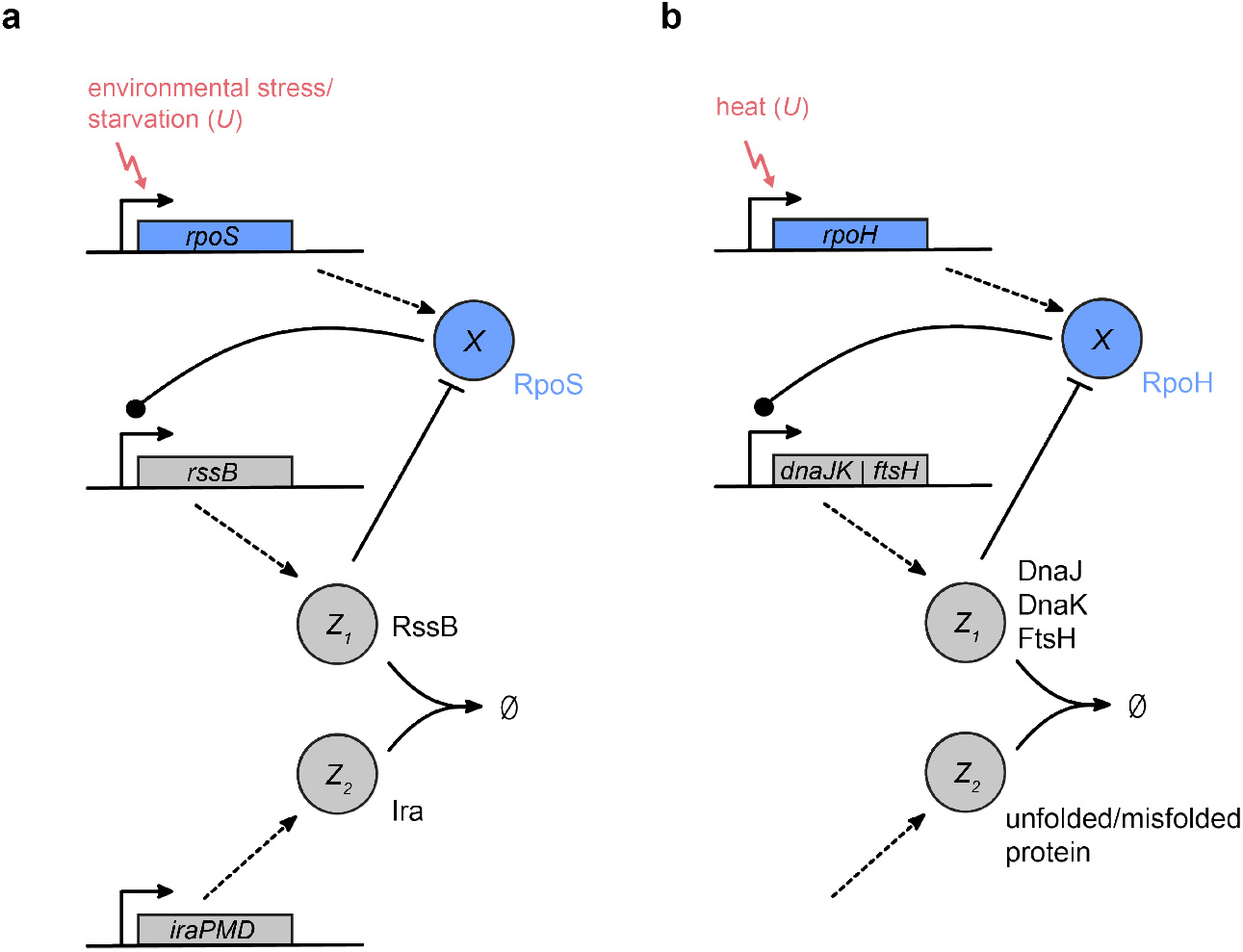
Examples of the Biomolecular Signal Differentiator-II motif in natural systems. Simplified schematics of BioSD-II topologies occurring as part of **a** the RpoS-mediated stress response and **b** the RpoH-mediated heat shock response in *Escherichia coli*. Corresponding components of BioSD-II are indicated.

#### Heat shock response – RpoH regulatory network

A second example for the BioSD-II motif was identified in the regulatory network of the sigma factor RpoH, which coordinates the heat shock response in *E. coli* (Fig. 6b) [37, 38]. Upon heat shock, cellular RpoH levels rise significantly above their low baseline concentrations (*b*), inducing the expression of several chaperones (e.g. DnaKJ and GroELS) and proteases (e.g. FtsH and Lon). DnaK and DnaJ can bind to RpoH and facilitate its degradation by FtsH [39, 40], thereby acting as *Z*_1_. Unfolded or misfolded proteins will sequester chaperones and proteases [40], thus increasing the stability of RpoH and fulfilling the function of *Z*_2_. In this network, the amount of *active* RpoH (as opposed to the total amount of RpoH) should be considered as *X*, since it has been found that the activity rather than the concentration of RpoH inside the cell drops during temperature downshifts [41].

## Discussion

In this study, we propose three biomolecular topologies that are able to act as highly accurate signal differentiators inside the cell. These designs provide guidance for building cellular devices capable of computing time derivatives of molecular signals. At the same time, they reveal concepts that are found in natural biological networks implementing differentiation and derivative feedback.

More specifically, we introduce three general biomolecular architectures BioSD-I, II and III. Their generality lies in the fact that they are represented by chemical reaction networks without being restricted by the biological identity of reactants and products and, by extension, the corresponding biological pathway. Important structural components of the BioSDs are a negative feedback loop created by a special process of excitation and inhibition between two species [42], an enzymatic degradation of zero-order kinetics (BioSD-I), an autocatalytic-like reaction (BioSD-III) and an antithetic-like motif based on annihilation [21, 25] (BioSD-II, BioSD-III). We theoretically analyze their features and show the conditions under which high performance can be guaranteed. Among others, important concepts such as stability, tunability and accuracy are discussed in detail.

Special emphasis is placed on the expected sensitivity of differentiators to input signals corrupted by high-frequency noise. We demonstrate that this issue can be resolved to a certain extent simply through suitable parameter tuning. Nevertheless, for cases in which stronger noise attenuation is needed, we present a structural modification that gives rise to three slightly different architectures, namely BioSD^*F*^-I, II and III, with enhanced capabilities. However, the price for this improvement is the addition of an extra biomolecular species, which implies an increase in structural complexity. Furthermore, we introduce performance metrics both for BioSDs (Supplementary Note 5) and BioSD^*F*^ s (Supplementary Note 6) based on which the circuit designer can assess the quality of signal differentiation and attenuation. These metrics take into account both the frequency content of the input signal and the reaction rates involved in the circuits, thus facilitating tuning according to the expected performance standards.

The ability to perform time differentiation is of central importance in various biological systems, contributing to stability and fast adaptation to changing conditions [6, 9, 43]. Owing to the generality of the presented topologies, we anticipate that the present study will facilitate the investigation of naturally occurring systems capable of derivative action. In this study we discuss the regulatory networks of two bacterial sigma factors, RpoS and RpoH, which play a central role in the response and adaptation to stress conditions and heat shock, respectively. Interestingly, these networks share structural characteristics with one of the proposed topologies, BioSD-II.

In addition, the motifs presented here provide building blocks that can be both implemented in stand-alone applications, such as speed biosensors, and also combined with existing biochemical control structures in a modular fashion, e.g. for building biomolecular PID controllers [18]. We describe potential designs for synthetic experimental implementation of all three BioSDs, which can be readily adapted depending on the nature of the system and available biological parts (Supplementary Note 8). To realize the antithetic motif in BioSD-II and III, we propose the use of a protease/protease inhibitor pair as an alternative to the previously described systems using sigma and anti-sigma factors [44] or sRNAs and mRNAs [45, 46]. To allow for greater flexibility in choosing the biomolecular species, we introduce the concept of auxiliary species, which is further analyzed in Supplementary Note 9.

The speed or higher derivatives of the output of a system offers important information about its properties. For an electromechanical system this is not difficult, but this has been a challenging question for biological systems. In this paper we provide an approach to gain access to this information, which will be invaluable for assessing and improving the performance of biological systems. We believe that our BioSD topologies will expand the tools available for understanding and engineering biological systems for robustness and reliability.

## Code availability

All numerical simulations were performed in MATLAB R2020 using the ODE solver ode23s except for those in Fig. 5 where the ODE solver ode113 was used. Simulation parameter values can be found in the figure captions. Initial conditions for the biomolecular species involved are considered zero except for BioSDs and BioSDs^*F*^ where the corresponding equilibria (“rest-positions”) are used (see Supplementary Note 1 and Supplementary Note 6).

## Acknowledgements

This work was supported by funding from the Engineering and Physical Sciences Research Council (EPSRC) [grant numbers EP/M002454/1 and EP/L016494/1] and the Biotechnology and Biological Sciences Research Council (BBSRC) [grant number BB/M011224/1]. C.C.M.S. is supported by the Clarendon Fund (Oxford University Press) and the Keble College De Breyne Scholarship. L.C. is supported by a Royal Society Research Professorship.

## Author contributions

E.A. conceived the theoretical designs and performed the mathematical analysis and computational simulations. C.C.M.S. and E.A. proposed experimental realizations. A.P. and L.C. designed and supervised the research. All authors contributed to the preparation and approved the submitted manuscript.

## Supplementary Information

### Supplementary Note 1: Equilibria and stability of Biomolecular Signal Differentiators

We assume that all biomolecular circuits in this study are represented by chemical reaction networks (CRNs) whose dynamics are described by the law of mass action unless otherwise stated. For the purposes of deterministic modelling, we consider input *U*(*t*) as a bounded, non-negative, continuous-time signal, the time derivatives of which exist and are also bounded. This is clearly aligned with the biological nature of *U*(*t*) which can correspond, for example, to the concentration of a biomolecular species.

#### Supplementary Note 1.1: Biomolecular Signal Differentiator-I

Biomolecular Signal Differentiator-I (BioSD-I) is described by the CRN:

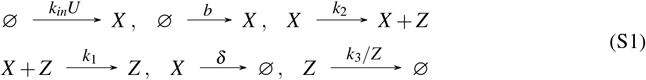

where *k_in_, b, k*_2_, *k*_1_, *k*_3_, 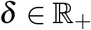. Note that the removal rate of *Z* is constant and equal to *k*_3_. This is possible if *Z* participates in an enzyme-catalyzed degradation process which is traditionally described by Michaelis-Menten kinetics. More precisely, the removal rate of *Z* is equal to

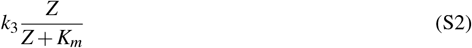

where 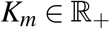 is the Michaelis-Menten constant. When the enzyme that catalyzes the degradation process is saturated by its substrate, we have:

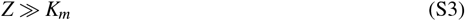

which entails, in effect, zero-order kinetics since Eq. (S2) becomes practically equal to *k*_3_.

The dynamics of CRN (S1) are given by the following system of Ordinary Differential Equations (ODEs):

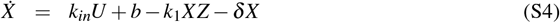

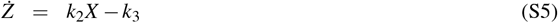

For any constant input *U*\*, a steady state (*X*\*, *Z*\*) of the system (S4), (S5) exists and is finite. By setting the time derivatives of this system to zero, we can obtain the following unique steady-state:

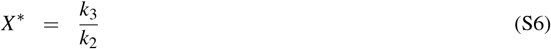

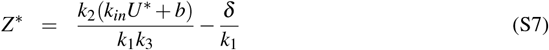

Clearly *X*\* is positive while, due to Eq. (S3), the same is true for *Z*\* (in fact: *Z*\* ≫ 0).

To study the local stability of the above equilibrium, we linearize (S4) – (S5) around (*X*\*, *Z*\*) for a constant input *U*\* to get:

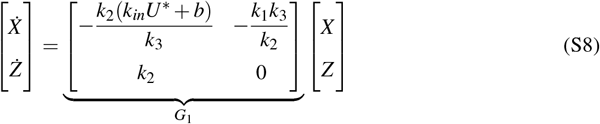

As far as the linear system (S8) is concerned, the steady state (*X*\*, *Z*\*) is exponentially stable since matrix *G*_1_ is Hurwitz. To prove this, we find the characteristic polynomial of *G*_1_ as:

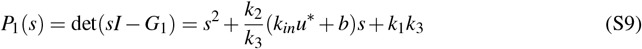

According to Routh-Hurwitz criterion, the second-order polynomial (S9) has both roots in the open left half plane if and only if both 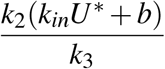 and *k*_1_*k*_3_ are positive, which is always true. Consequently, (*X*\*, *Z*\*) is a positive locally exponentially stable steady state for the nonlinear system (S4), (S5).

Following the same procedure, we next analyze BioSD-II and BioSD-III.

#### Supplementary Note 1.2: Biomolecular Signal Differentiator-II

The CRN that corresponds to Biomolecular Signal Differentiator-II (BioSD-II) is:

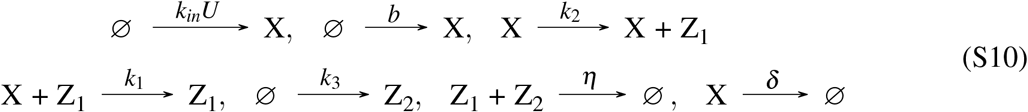

where *k_in_, b, k*_2_, *k*_1_, *δ*, 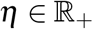.

The dynamics of CRN (S10) are described by the set of ODEs:

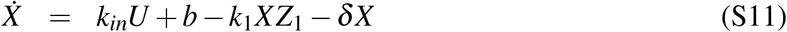

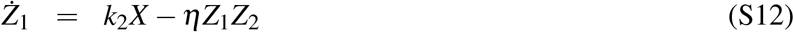

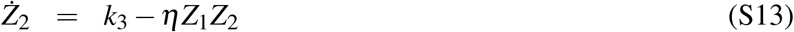

For any constant input *U*\*, provided that:

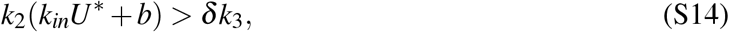

we have a unique positive (finite) steady state:

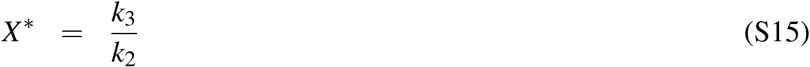

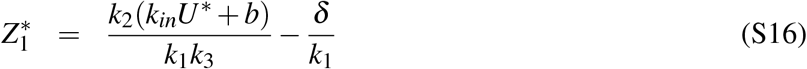

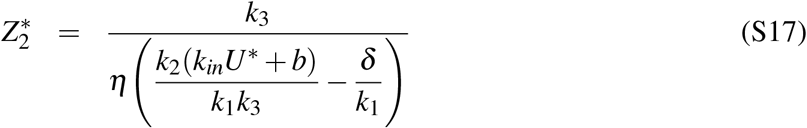

We now linearize (S11) – (S13) around the fixed point defined by (S15) - (S17) to obtain:

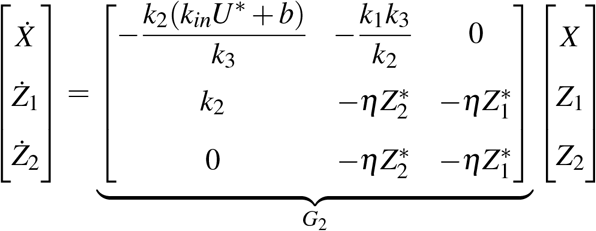

The characteristic polynomial of *G*_2_ is:

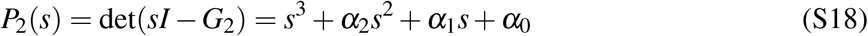

where

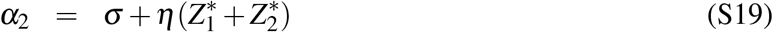

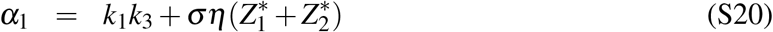

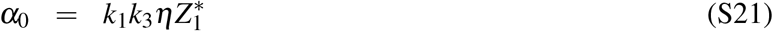

and

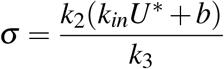

The polynomial (S18) has all roots in the open-half plane if and only if *α*_2_, *α*_0_ are positive and *α*_2_*α*_1_ > *α*_0_ (Routh-Hurwitz criterion). Indeed:

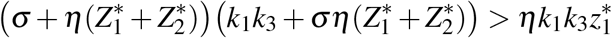

or

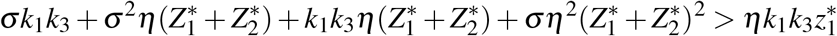

or

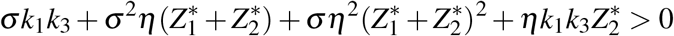

which is always true since all the quantities involved are positive. Therefore, 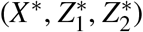 is a positive locally exponentially stable steady state (*G*_2_ is Hurwitz) for the nonlinear system (S11) - (S13).

Note that outside the parameter regime defined by Eq. (S14) BioSD-II is unable to reach equilibrium. In particular, assuming non-negative initial conditions for the system (S11) – (S13) (which is always the case because the variables involved represent biomolecular concentrations) the states of the latter remain always non-negative (as expected from mass action kinetics). Indeed, when *X* = 0, Eq. (S11) implies *Ẋ* = *k_in_U* + *b* > 0. Furthermore, when *Z*_1_ = 0, Eq. (S12) results in *Ż*_1_ = *k*_2_*X* ≥ 0 and, finally, when *Z*_2_ = 0, Eq. (S13) imposes *Ż*_2_ = *k*_3_ > 0. However, outside the parameter regime in question, one of the following must hold: *k*_2_(*k_in_U*\* + *b*) < *δk*_3_ or *k*_2_(*k_in_U*\* + *b*) = *δk*_3_. In the first scenario, it is apparent from Eqs. (S16), (S17) that the steady state of *Z*_1_, *Z*_2_ becomes negative while in the second case Eq. (S17) indicates that *Z*_2_ tends to infinity - thus, BioSD-II cannot approach a finite steady state.

#### Supplementary Note 1.3: Biomolecular Signal Differentiator-III

Biomolecular Signal Differentiator-III (BioSD-III) is represented by the CRN:

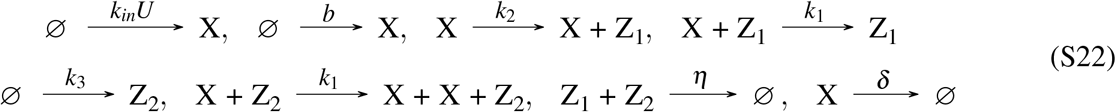

where *k_in_, b, k*_2_, *k*_1_, *δ*, 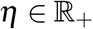.

The corresponding ODE model describing the dynamics is:

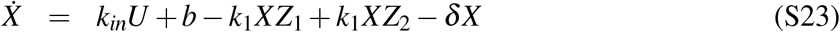

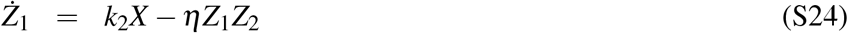

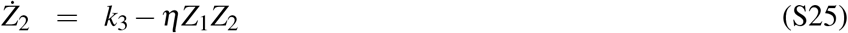

For any constant input *U*\*, we have a unique positive steady state (providing that it exists and is finite):

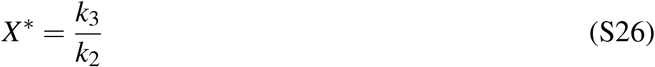

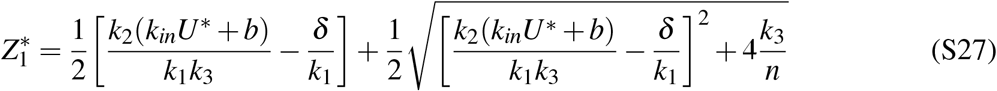

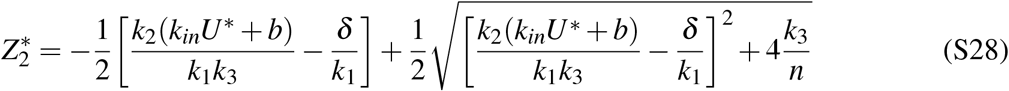

Linearizing the system (S23)-(S25) around its steady state (S26)-(S28) yields:

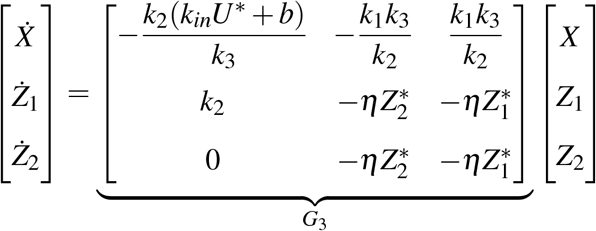

The characteristic polynomial of *G*_3_ is:

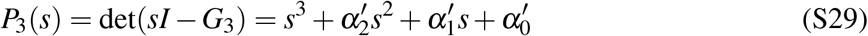

where 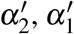 are identical to *α*_2_ (Eq. S19), *α*_1_ (Eq. S20), respectively and:

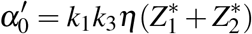

In order to show that *G*_3_ is Hurwitz we need to verify that 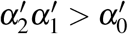 (from Routh-Hurwitz criterion).

This inequality is satisfied because:

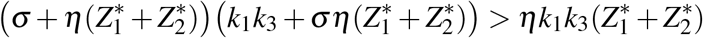

or

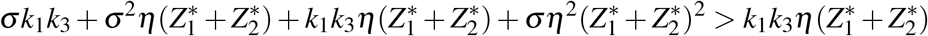

or

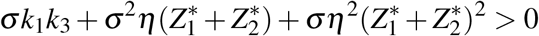

which is always true as a sum of positive quantities. Hence, 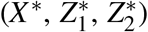 is a positive locally exponentially stable steady state for the nonlinear system (S23) - (S25).

### Supplementary Note 2: The notion of strong rate of annihilation between *Z*_1_, *Z*_2_ (large *η*) in Biomolecular Signal Differentiator-II

BioSD-II involves the following annihilation reaction (CRN (S10)):

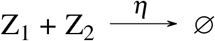

This reaction describes a process where species *Z*_1_, *Z*_2_ bind to each other irreversibly to form a product which can be considered as biologically inactive. In other words, this product does not participate in any of the reactions in BioSD-II. Here we demonstrate that near the equilibrium the effect of species *Z*_2_ can be considered negligible if the formation rate, *η*, of the product in question is sufficiently high. As we show in the next section, this is a requirement in order for BioSD-II to exhibit accurate signal differentiation.

By adopting the coordinate transformations: *u* = *U* – *U*\*, *x* = *X* – *X*\*, 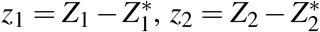 which denote small perturbations around 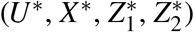, we obtain through linearization of (S11) – (S13):

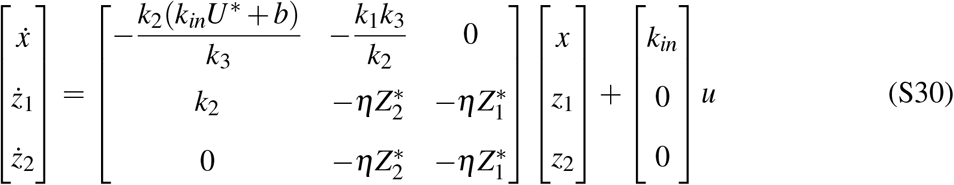

We now introduce the nondimensional variables:

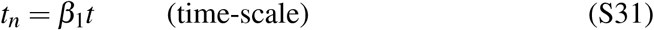

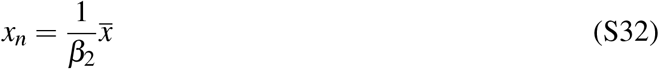

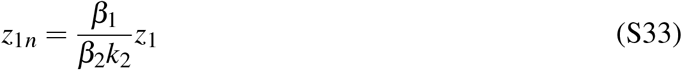

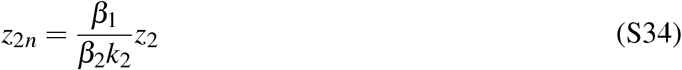

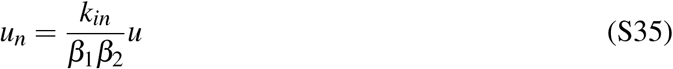

where

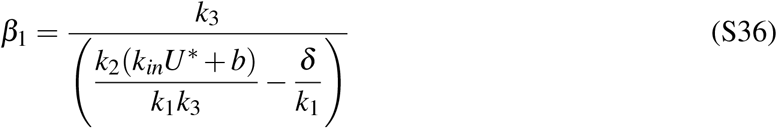

and *β*_2_ is an arbitrary scaling parameter that carries the same units as *x_n_*. In addition, we introduce the nondimensional parameters:

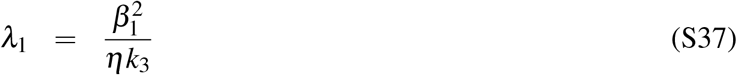

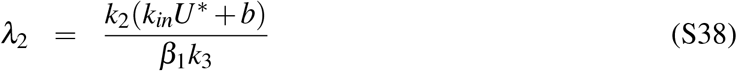

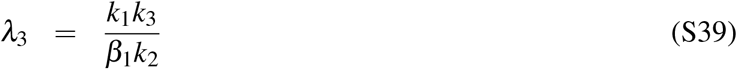

By substituting Eqs. (S31)–(S39) into the model (S30), we obtain:

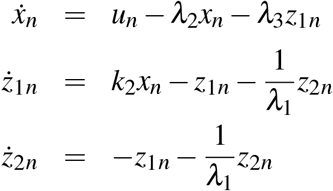

or

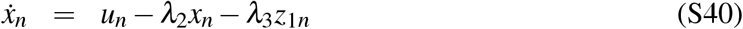

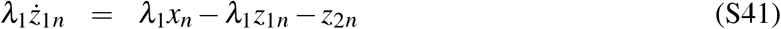

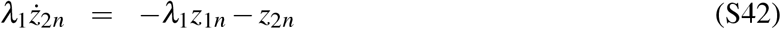

According to Eq. (S37), *λ*_1_ → 0 as *η* → ∞. This means that we can make *λ*_1_ negligible by choosing a large value for *η*:

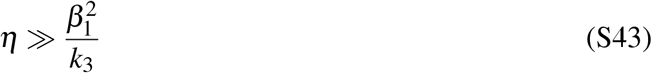

Assuming now that *λ*_1_ is zero the ODE model (S40)–(S42) becomes:

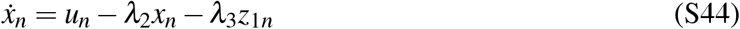

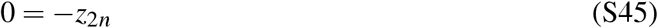

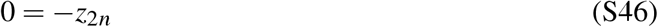

Combining Eqs. (S45), (S46) with Eq. (S34), we get:

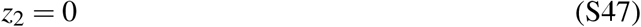

From Eqs. (S34), (S47) and taking into account that 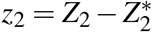 we obtain 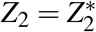. Moreover, the combination of (S16), (S36) and (S37) yields:

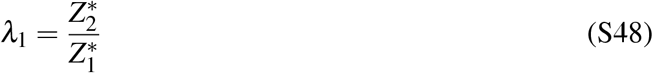

Assuming that 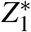 is finite and *λ*_1_ is zero we have 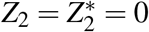.

As a consequence, if the condition (S43) is true (*λ*_1_ is insignificant) then the deviation, *z*_2_, of species *Z*_2_ from its steady state, 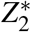, can be considered effectively zero while the latter is negligible compared to 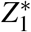. Furthermore, the case for which the parameter *η* satisfies Eq. (S43) will be referred to as “sufficiently large *η*”.

### Supplementary Note 3: Signal differentiation

Here we prove that, near their equilibria, BioSD networks are capable of signal differentiation. We begin with BioSD-I whose dynamics close to its steady state are derived via linearization of Eqs. (S4), (S5) as:

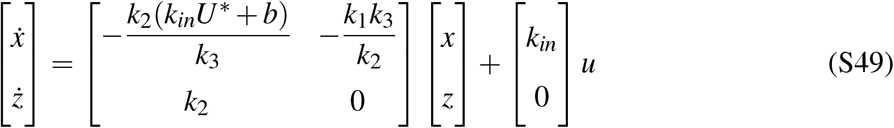

assuming the coordinate transformations: *u* = *U* – *U*\*, *x* = *X* – *X*\*, *z* = *Z* – *Z*\* which represent small perturbations around (*U*\*, *X*\*, *Z*\*). We next consider the nondimensional variables:

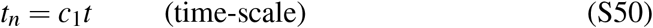

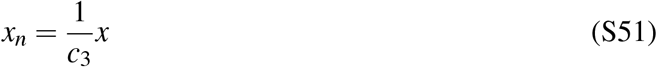

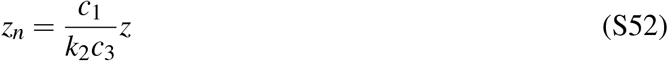

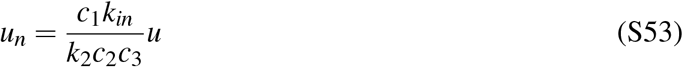

where

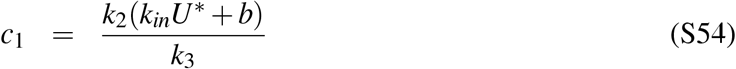

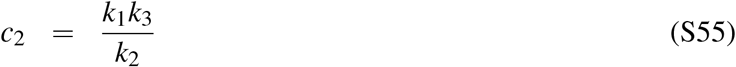

and *c*_3_ is an arbitrary scaling parameter that carries the same units as *x_n_*. We also introduce the non-dimensional parameter:

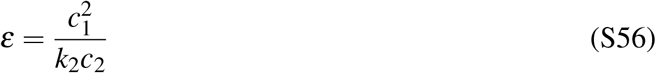

Substituting Eqs. (S50)–(S56) into the system (S49) results in:

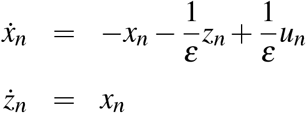

or

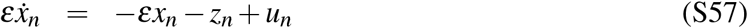

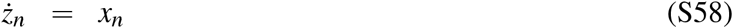

We now turn our focus to the scenario where *ε* is small. This can be achieved in practice by selecting model parameters such that *ε* ≪ 1. Taking into account Eqs. (S54), (S55) and (S56) we have:

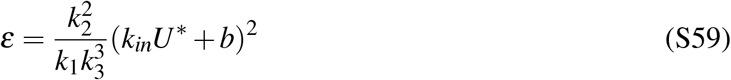

and, consequently, the aforementioned condition can be reformulated as:

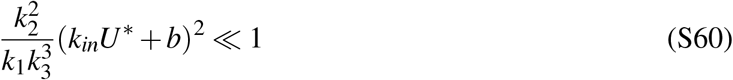

We now study the ODE model (S57), (S58) under the assumption that *ε* is equal to zero. Thus, we have:

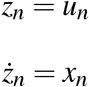

or

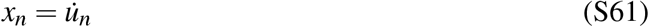

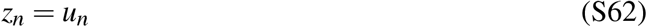

Eq. (S61) becomes through Eqs. (S50), (S51), (S53), (S55):

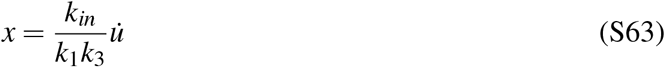

By recalling Eq. (S6) and our initial coordinate transformations, this relationship can be rewritten as:

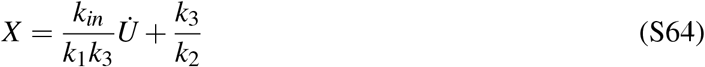

The above result states that BioSD-I produces an output proportional to the derivative of the input and shifted by a fixed amount equal to 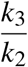. The latter amount can be viewed as the zero-level ‘bias’ on which the derivative of the input is evaluated. This “uplifting” is undoubtedly necessary in order for negative values of the derivative to be represented by a biomolecular species.

We now construct the following second-order differential equation using Eqs. (S49), (S54) and (S55):

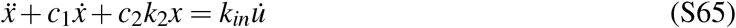

We see immediately that if:

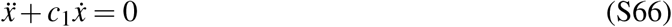

then we arrive at Eq. (S64) or, in other words, we have identical derivative action as before. Therefore, we find the general solution of (S66) which is:

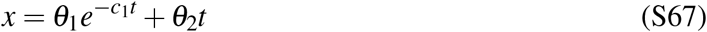

where *θ*_1_, *θ*_2_ are arbitrary constants. Subsequently, from Eqs. (S63), (S67) we get:

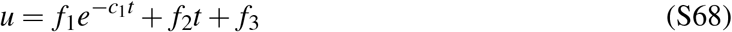

where *f*_1_, *f*_2_, *f*_3_ are arbitrary constants 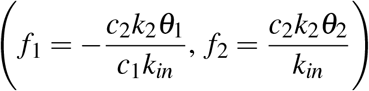.

As a consequence, for any input signal that is described or can be approximated by (S68), the biological differentiator in question does not require special handling of the system parameters in order to satisfy the constraint (S60) since it works satisfactorily even in the case where *ε* is far away from zero.

Following the same procedure, we study the local dynamics of BioSD-III by linearizing Eqs. (S23)–(S25):

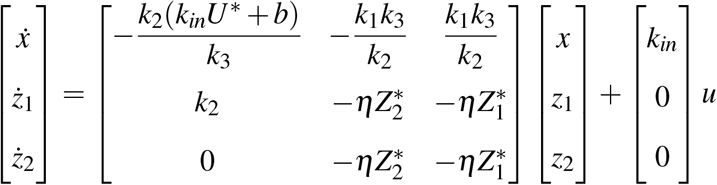

where the variables *u* = *U* – *U*\*, *x* = *X* – *X*\*, 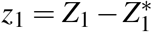, 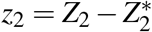 refer to small perturbations around the equilibrium 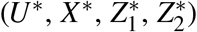. Introducing the linear transformation *g* = *z*_1_ – *z*_2_ results in the following mathematically equivalent system:

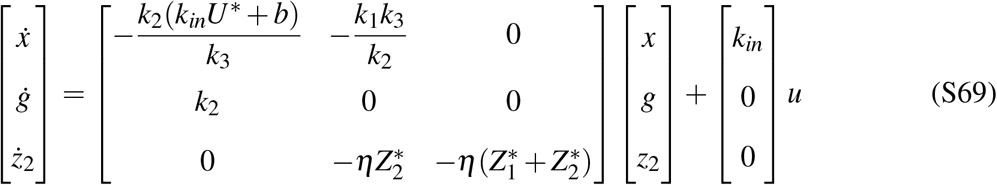

We notice that the dynamics of *x* and *g* of the system (S69) are identical to that of *x* and *z* of the system (S49), respectively. Hence, the output, *x*, of BioSD-III behaves in the exact same way as the one of previously analyzed BioSD-I.

Subsequently, we recall the dynamics of BioSD-II near its equilibrium (S30) from the preceding section. It is evident that using the linear transformation *g* = *z*_1_ – *z*_2_ again and assuming a sufficiently large *η* (condition (S43) is satisfied), the dynamics of *x* and *g* in BioSD-II are described by Eq. (S49), namely the dynamics of BioSD-I. By extension, the output behaviour of these two circuits is identical.

Finally, it is important to emphasize that the nonlinear systems (S4)-(S5), (S11)-(S13), (S23)-(S25) representing BioSD-I, BioSD-II, BioSD-III, respectively, are locally input-to-state stable. This is guaranteed by the fact that all the three aforementioned systems are differentiable in time and their equilibrium is locally asymptotically (and more specifically exponentially) stable, as shown in Supplementary Note 1.

### Supplementary Note 4: Accuracy of signal differentiation and its dependence on model parameters

As previously pointed out, a requirement for reliable signal differentiation is selection of such model parameters that the nondimensional parameter *ε* (given by Eq. (S60)) is sufficiently small. Here, we are interested in the impact of *ε* on the accuracy with which BioSD devices are able to differentiate the input signals applied. In particular, we investigate how the deviation of their behaviour from that of an ideal differentiator changes in the long term as *ε* decreases.

The input/output relation of an ideal differentiator is described in the Laplace domain by the transfer function:

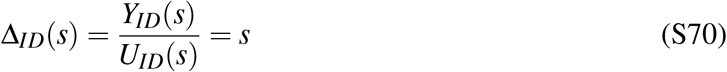

where *Y_ID_*(*s*) and *U_ID_*(*s*) refer to the Laplace transform of the output and input of the ideal differentiator, respectively. In addition, *s* = *σ* + *jω* is the frequency domain variable with *σ*, 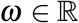 and *j* is the imaginary unit number 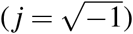.

Based on the analysis in Supplementary Note 3, we can derive a transfer function for BioSD differentiators from Eqs. (S57)), (S58)) as:

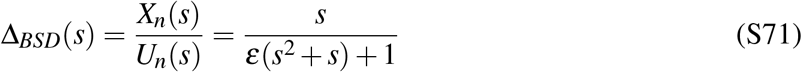

where *X_n_*(*s*) and *U_n_*(*s*) represent the Laplace transform of the output *x_n_* and input *u_n_* of BioSD networks.

Clearly the transfer functions (S70) and (S74) coincide when *ε* is zero. We now consider a positive *ε* in some neighbourhood of zero. Focusing on the frequency response (we set *σ* = 0 and essentially use Fourier transform), we have for an arbitrary (finite) frequency *ω*:

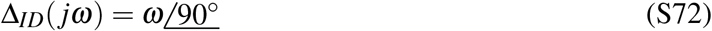

and

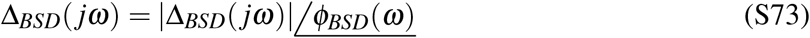

where:

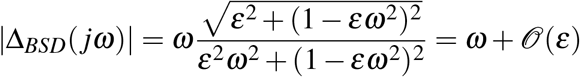

and

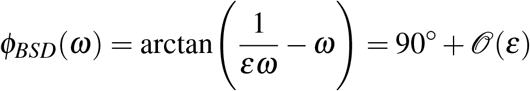

Furthermore, we calculate:

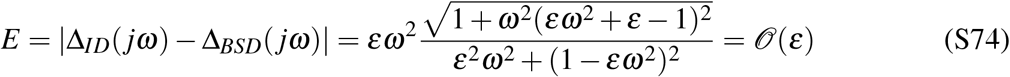

The above analysis reveals that in the long term, compared to an ideal differentiator, signal differentiation via BioSD modules implies a magnitude and phase shift of only 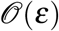. In parallel, the absolute value of the difference between the ideal transfer function and the actual one decreases proportionally with *ε*. Thus, as we decrease *ε*, the output of BioSD networks approaches the ideal derivative of the input and, by extension, the accuracy of signal differentiation provided increases.

### Supplementary Note 5: Changes in the behaviour of Biomolecular Signal Differentiators based on the frequency content of input signals

As shown in Supplementary Figure 1, there are two large areas of input frequencies, one with “lower frequencies” and one with “higher frequencies”, over which BioSD networks (*ε* > 0) can effectively perform signal differentiation and signal attenuation, respectively. It is evident that, as *ε* increases, the latter area expands towards lower frequencies and, thus, the former area shrinks. We can also detect the existence of a narrow frequency band between the two where BioSD circuits are not able to differentiate or attenuate input signals with the expected accuracy.

On the other hand, in the ideal case of *ε* = 0, perfect differentiation of input signals takes place regardless of their frequency content. Although a behaviour like this may seem desirable, in practice it can be catastrophic since it leads to substantial amplification of high frequency signal components (including noise from the cellular environment). As a consequence, the “imperfection” of BioSDs turns out to be a saving feature of great significance.

We now recall Eqs. (S72), (S73) that describe the frequency response from the input we want to differentiate to the output we measure regarding an ideal differentiator and a BioSD, respectively. As already mentioned, it is obvious from these relationships that for input signals which remain constant over time (*ω* = 0), the two systems operate in the exact same way. On the contrary, for time-varying input signals (*ω* > 0), considerable deviations in their behaviour may appear, as pointed out above. To get a sense of how close the function of a BioSD device is to the ideal function for an arbitrary input frequency, we introduce:

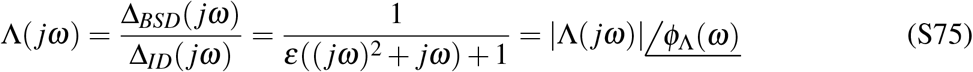

where

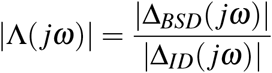

and

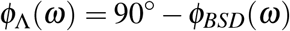

When a BioSD network operates exactly as an ideal differentiator, then 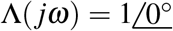, whereas if |Λ(*jω*)| = 0 perfect signal attenuation is achieved. Having these extreme situations in mind and taking into account the expected performance standards, the circuit designer can use Eq. (S75) as a metric in order to select a suitable *ε* and, by extension, an appropriate combination of model parameters. Finally, Supplementary Figure 2 illustrates for different values of *ε* how the performance metric in question changes based on the input frequencies.

### Supplementary Note 6: An alternative version of Biomolecular Signal Differentiators

Here we analyze a slightly different version of the previously studied BioSD networks which we call Biomolecular Signal Differentiators^*F*^ (BioSDs^*F*^) that include an additional biomolecular species, *Z*_3_. In particular, we describe the following three biomolecular topologies:

- **BioSD^*F*^-I** We have the CRN:

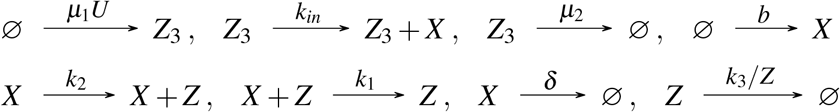

where *μ*_1_, *μ*_2_, *k_in_*, *b*, *k*_2_, *k*_1_, *δ*, 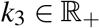. The corresponding ODE model is:

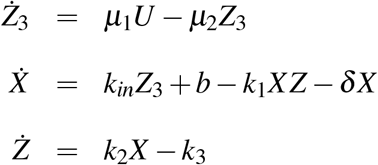
- **BioSD^*F*^-II** We have the CRN:

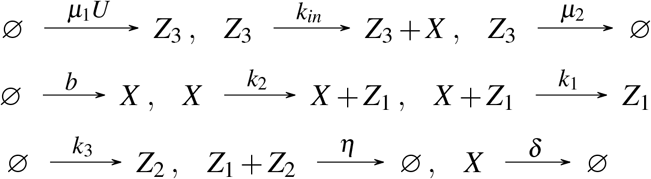

where *μ*_1_, *μ*_2_, *k_in_, b, k*_2_, *k*_1_, *δ*, 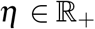. We assume that the parameter rate *η* is sufficiently large (see Supplementary Note 2). The corresponding ODE model is:

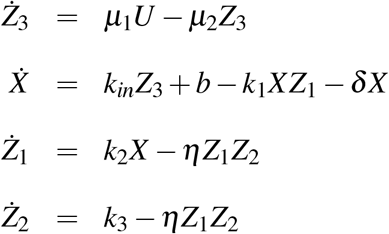
- **BioSD^*F*^-III** We have the CRN:

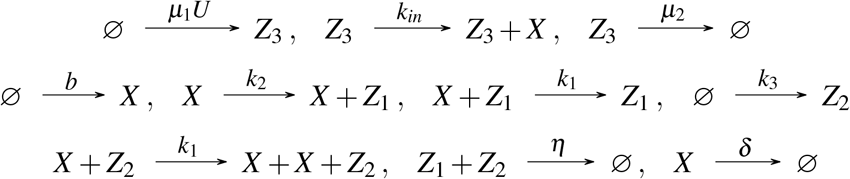

where *μ*_1_, *μ*_2_, *k_in_*, *b, k*_2_, *k*_1_, *δ*, 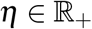. The corresponding ODE model is:

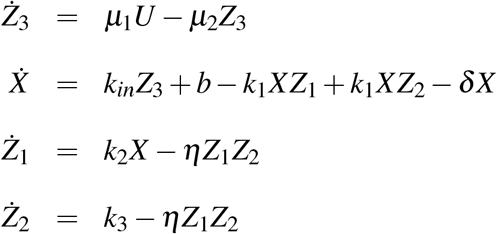

It is evident that each of the above circuits can be seen as the interconnection of two subsystems. More specifically, we have the linear subsystem (the first equation in each of above ODE models):

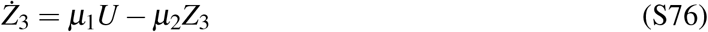

which receives the signal *U* we want to differentiate as input and produces an output *Z*_3_. This is, in turn, applied as input to a second subsystem whose output is *X*. While the first subsystem is the same in all BioSD^*F*^ topologies, the second one differs. In fact, the latter is identical to BioSD-I, BioSD-II, BioSD-III (see previous sections) for BioSD^*F*^-I, BioSD^*F*^-II, BioSD^*F*^-III, respectively, with the only difference lying in the input, which is now *Z*_3_ (instead of *U* as before).

For a constant input *U*\* the first subsystem (S76) has a unique positive steady state (assuming it exists and is finite):

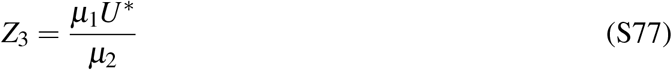

Since Eq. (S76) is linear and (*μ*_2_) is always positive then Eq. (S77) constitutes a globally exponentially stable equilibrium point. This also guarantees global input-to-state stability.

We now concentrate on the local behaviour of BioSD^*F*^ modules and, consequently, we consider the coordinate transformations: *u* = *U* – *U*\*, *x* = *X* – *X*\*, *z* = *Z* – *Z*\*, 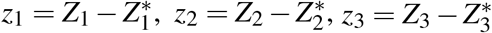 denoting small perturbations around the corresponding equilibria of BioSD^*F*^ networks - 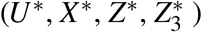 for BioSD^*F*^-I and 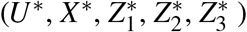 for BioSD^*F*^-II, BioSD^*F*^-III (the steady state of the last two networks do not necessarily coincide).

First, we study Eq. (S76) separately. In the Laplace domain, we have:

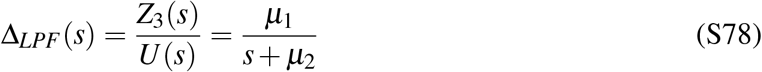

where *Z*_3_(*s*), *U*(*s*) are the Laplace transform of *z*_3_, *u*, respectively. Focusing on the frequency response, we get:

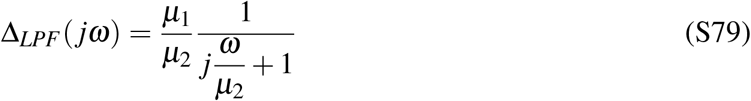

This is a transfer function of a first-order low-pass filter which is capable of preserving low-frequency signals and rejecting high-frequency signals. Indeed, the magnitude and the phase of the system in question are given by:

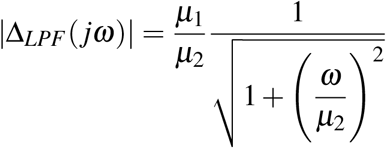

and

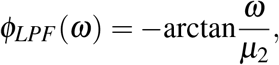

respectively.

We can easily see that in practice, when *ω* ≪ *μ*_2_, there is a constant input/output gain 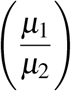 and no phase lag. On the other hand, for 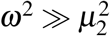 strong attenuation takes place. The general behaviour of the filter can be easily understood through the Bode diagram in Supplementary Figure 3.

We now consider a BioSD^*F*^ design which can be described by the transfer function of the series connection of the previously studied filter and a BioSD design (as already outlined in Supplementary Note 4, all three BioSD circuits are described by the same transfer function), i.e.:

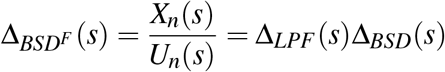

or

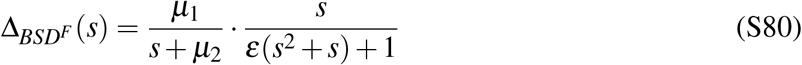

where 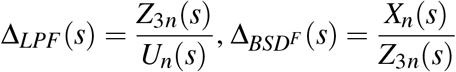 with *Z*_3*n*_(*s*) = *pZ*_3_(*s*) and 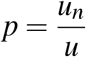 (Supplementary Note 3).

For 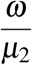, *ε* → 0 Eq. (S80) becomes:

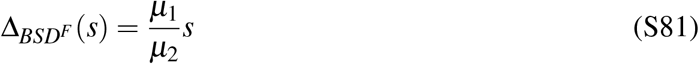

which implies in the time domain (recall Supplementary Note 3):

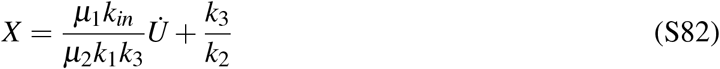

Hence, in this case, BioSD^*F*^ networks work in a similar manner as BioSDs (Eq. (S64)) with respect to signal differentiation. The only difference appears in the output gain by which the derivative of the input is multiplied. Besides *k_in_, k*_1_, *k*_3_, this gain is now determined by *μ*_1_, *μ*_2_ as well, thus providing increased tunability.

Next, we shift our focus to the more general scenario where we allow *ε* to move in some neighbourhood of zero and 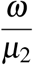 to vary from zero to infinity. We can easily evaluate the performance of a BioSD^*F*^ differentiator by taking advantage of our initial decomposition. In particular, we can predict the behaviour of the differentiator from the behaviour of its individual components, namely a low-pass filter (analyzed earlier in this section) and a BioSD device (see the preceding sections and especially Supplementary Note 4 and 5). Indeed, exploiting frequency response analysis for performance assessment in the long term, we have for an arbitrary input frequency *ω*:

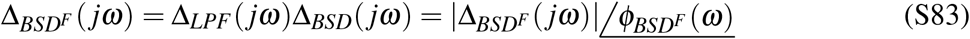

where

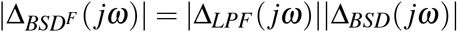

and

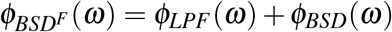

In addition, following the notion of Supplementary Note 5, we can introduce the (normalized) performance metric:

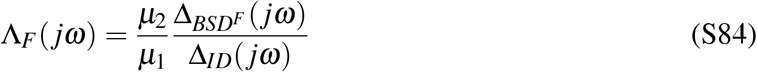

where

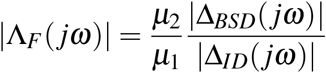

and

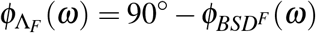

Perfect signal differentiation and attenuation happen when 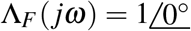 and |Λ_*F*_(*jω*)| = 0, respectively.

Consequently, for a given *ε*, BioSD^*F*^ circuits are characterized by an enhanced capability of high-frequency signal attenuation compared to BioSD ones. In fact, as demonstrated in Supplementary Figure 4, we can extend the frequency band where strong signal attenuation is carried out by appropriately tuning the filter module. In other words, we can adjust the bandwidth of the filter as desired through the parameter rate *μ*_2_. The price we pay for this significant improvement is the increase in structural complexity due to the addition of the species *Z*_3_ via which the additional filtering is accomplished. Finally, note that accompanying a differentiator with a low-pass filter is a widely used strategy in traditional engineering in order to deal with high-frequency measurement noise.

### Supplementary Note 7: Numerical simulations

In this section we present additional simulations for further computational validation of our theoretical results in section **Sensing the response speed of biomolecular networks** in the main text. In particular, we numerically investigate the response of BioSD-II and BioSD-III to the inputs shown in Fig. 2c and Fig. 3c. As can be seen from Supplementary Figure 5 and Supplementary Figure 6, the behaviour of both BioSD-II and BioSD-III is identical to that of BioSD-I depicted in the main text. As a result, the conclusions drawn with respect to the latter circuit are valid for the other designs as well.

### Supplementary Note 8: Guidelines for experimental implementation of Biomolecular Signal Differentiators

In the main text we have focused on naturally occurring examples of the BioSD-II topology. Here, we discuss how *synthetic* BioSD circuits can be designed and implemented inside a living cell. In particular, we propose experimental implementations of the described topologies in *Escherichia coli* (Supplementary Figure 7).

Inducible expression of species *X* can be achieved from any well-characterized promoter, such as the IPTG-inducible *P_lac_*. Leakiness of the lac promoter will ensure nonzero expression levels (*b*) even in the absence of inducer. Alternatively, if higher baseline expression levels are required, *X* could additionally be expressed from a weak constitutive promoter.

To minimize undesirable interference with other cellular processes, *X* should be an orthogonal sigma factor, such as σ^*F*^ from *Bacillus subtilis* [1]. A translational fusion of *X* to GFP will allow for easy tracking of the system output. *σ^F^* will then induce expression of a Lon protease (*Z* in BioSD-I, *Z*_1_ in BioSD-II and III) from its cognate promoter P_*F*1_. In this case, a Lon^−^ strain of *E. coli* would be used to avoid interference of naturally present Lon protease. Addition of a degradation tag to *σ^F^* will target it for degradation by the Lon protease. To approximate zeroth-order degradation, an *ssrA* tag will be fused to the Lon protease as previously described in [2, 3].

For BioSD-II, we additionally introduce constitutive expression of the protease inhibitor PinA from phage T4 (*Z*_2_), which has been shown to specifically inhibit the Lon protease in *E. coli* with high affinity [4]. A synthetic promoter from the BioBrick collection [5] may be used to achieve the desired expression level of *Z*_2_. Ideally, an orthogonal Lon protease should be used (e.g. Lon protease from *Mesoplasma florum* [6]) to prevent cross-talk with other cellular proteins. However, since the interaction of PinA with proteases has been characterized only in *E. coli* so far, we have suggested use of the *E. coli* Lon protease.

Due to the number of required interactions in BioSD-III, it will likely be necessary to introduce auxiliary species for *X, Z*_1_ and *Z*_2_, which we refer to as *X_aux_, Z*_1,*aux*_ and *Z*_2,*aux*_, respectively. These auxiliary species would ideally have identical behaviour to the main species *X, Z*_1_ and *Z*_2_, even though simulations indicate that completely identical behaviour is not required (see Supplementary Note 9).

One option is to augment the design for BioSD-II with the Hrp system from *Pseudomonas syringae*, which has previously been implemented in synthetic biology studies [7]. HrpR (*X_aux_*) fused to a fluorescent tag (mCherry) is expressed from P_*lac*_ together with *σ^F^*, and HrpS (*Z*_2,*aux*_) is expressed in an operon with PinA. HrpR and HrpS are both required to induce additional production of *σ^F^* and HrpR from P_*hrpL*_. At the same time, HrpV (*Z*_1,*aux*_) binds HrpS, rendering it inactive.

Finally, the structural addition required for BioSD^*F*^ can be implemented by, for example, expressing *X* from a T7 promoter and expressing T7 RNA polymerase (*Z*_3_) from a separate inducible promoter.

### Supplementary Note 9: Analysis of the experimental topology of Biomolecular Signal Differentiator-III

Here we further analyze the proposed synthetic design of BioSD-III, the behaviour of which may be more complicated due to the use of three auxiliary species (see Supplementary Note 8).

The biomolecular topology shown in Supplementary Figure 7c can be described by the following set of ODEs:

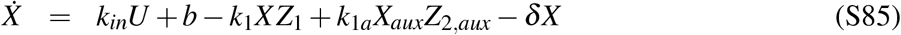

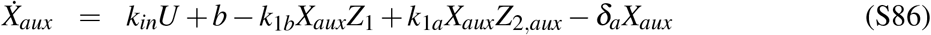

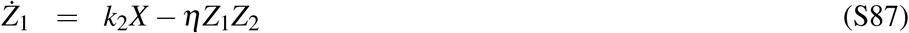

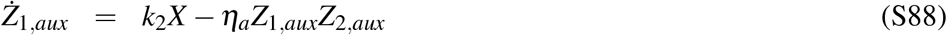

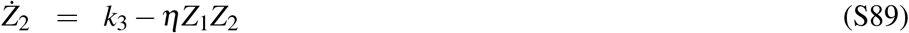

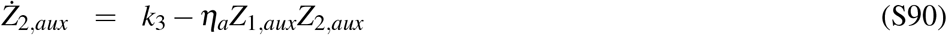

where *k_in_, b, k*_2_, *k*_1_, *k*_1,*a*_, *k*_1,*b*_, *δ, δ_a_, η*, 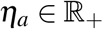.

In order for the behaviour of *X* (measured output species) in the system (S85)-(S90) to perfectly match the one of *X* in the model (S23)-(S25), we need: *k*_1_ = *k*_1*a*_ = *k*_1*b*_, *δ* = *δ_a_* and *η* = *η_a_*. Nevertheless, non-satisfaction of the aforementioned conditions does not necessarily entail considerable loss of accuracy regarding signal differentiation (Supplementary Figure 8).

**Supplementary Figure 1:**
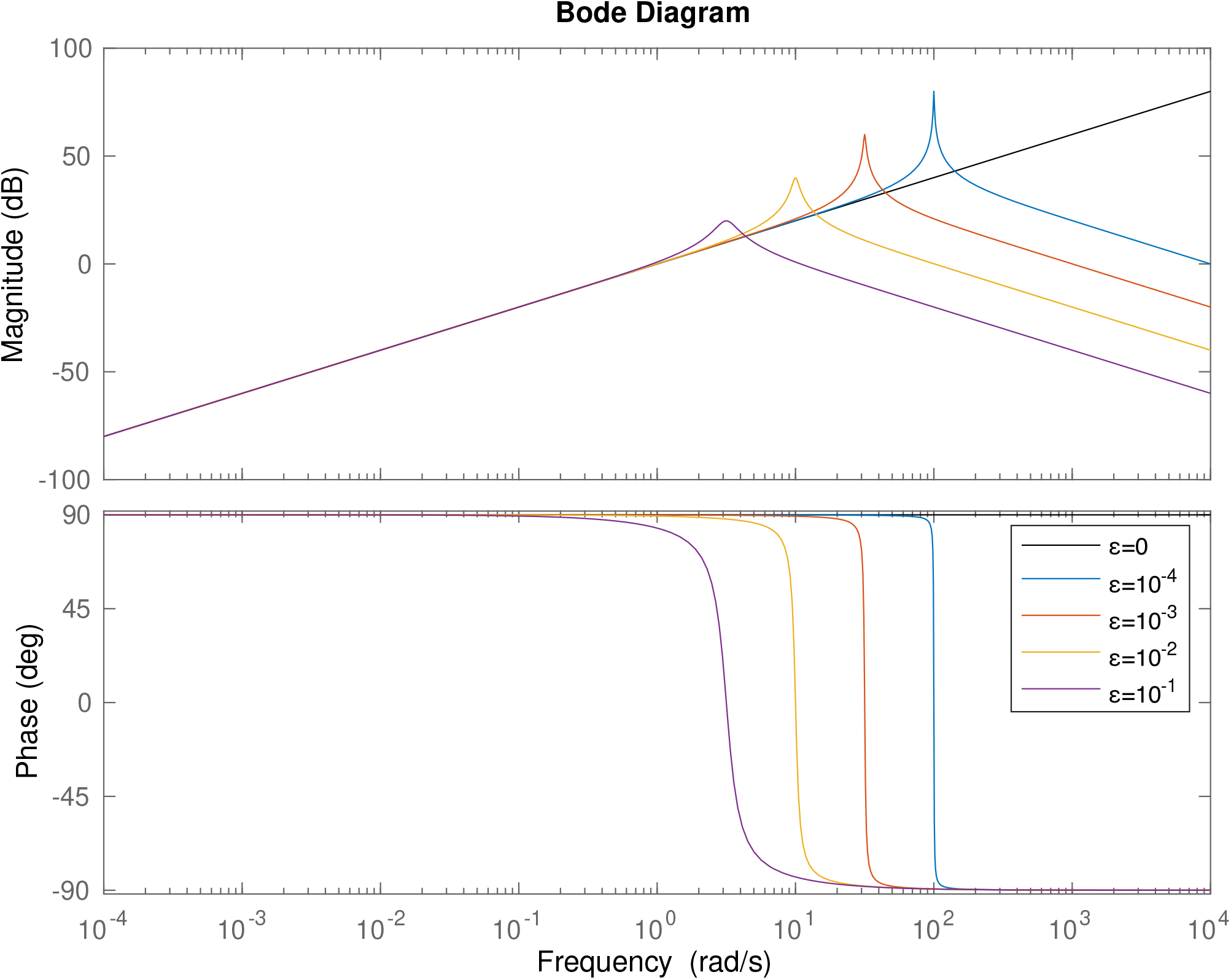
Frequency response analysis of Biomolecular Signal Differentiators. Bode plot of a BioSD differentiator (Eq. (S73)). The magnitude and the phase of its transfer function are depicted for different values of *ε* via distinct colours. The case of *ε* = 0 represents the behaviour of an ideal differentiator.

**Supplementary Figure 2:**
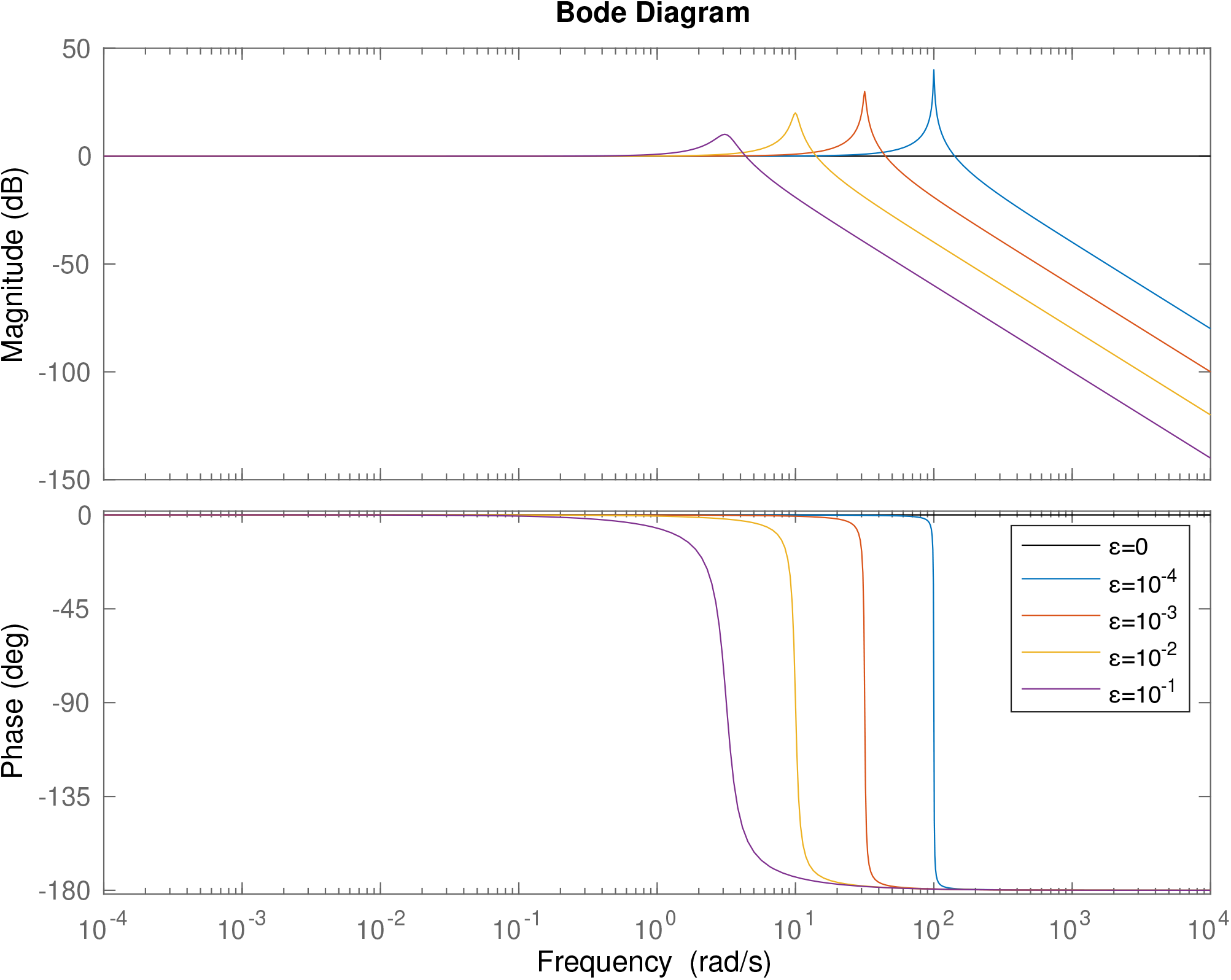
A performance metric in the frequency domain. Bode plot of the metric (S75) through which the accuracy of signal differentiation and signal attenuation regarding a BioSD can be easily assessed. Different colours represent the magnitude and the phase of the corresponding transfer function for different values of *ε*. The case of *ε* = 0 refers to an ideal differentiator.

**Supplementary Figure 3:**
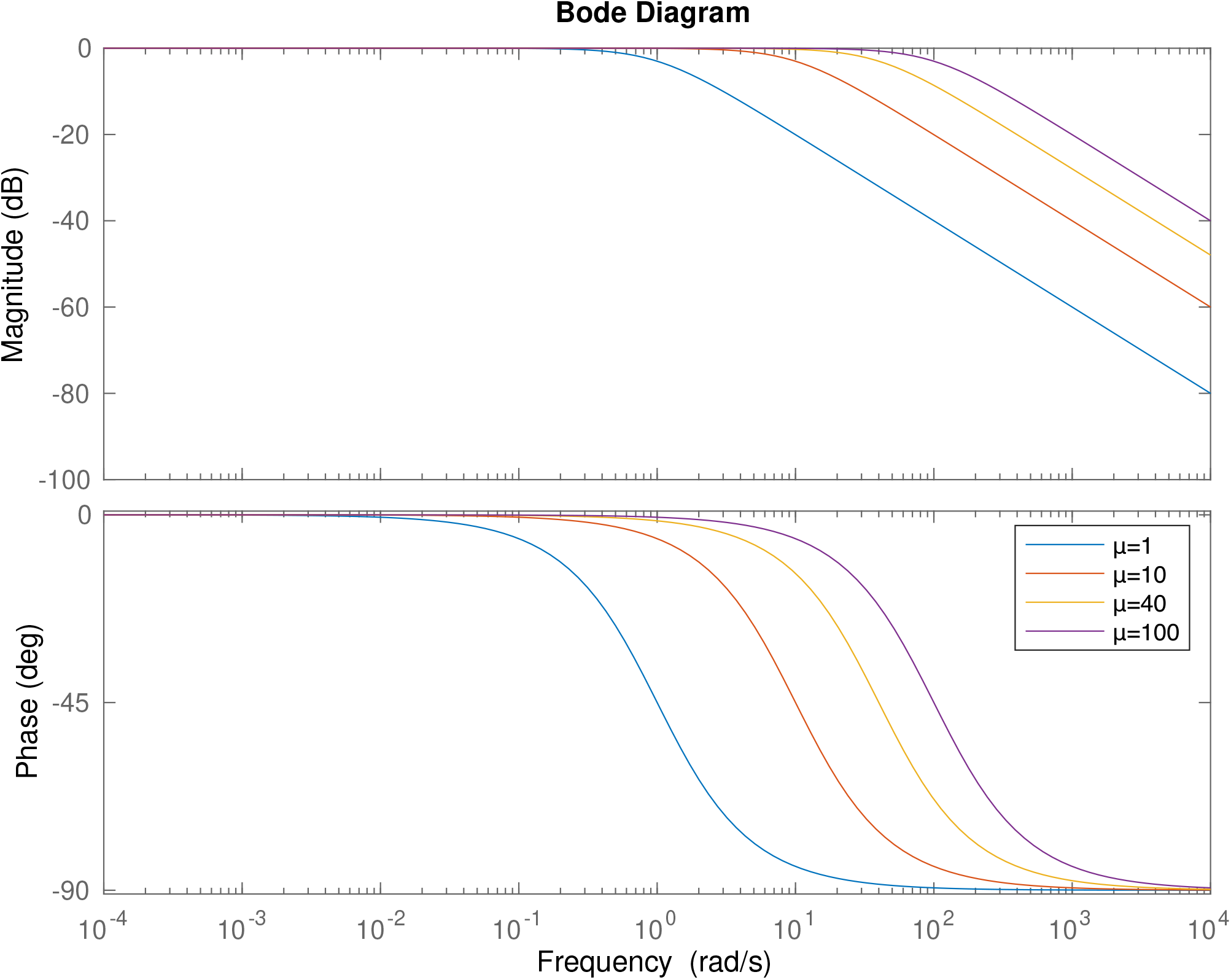
Frequency response analysis of the subsystem that receives the input signal *U*. Bode diagram of the filter module described by Eq. (S79). The magnitude and and the phase lag of its frequency response for different values of *μ* are shown in different colours where *μ* = *μ*_1_ = *μ*_2_.

**Supplementary Figure 4:**
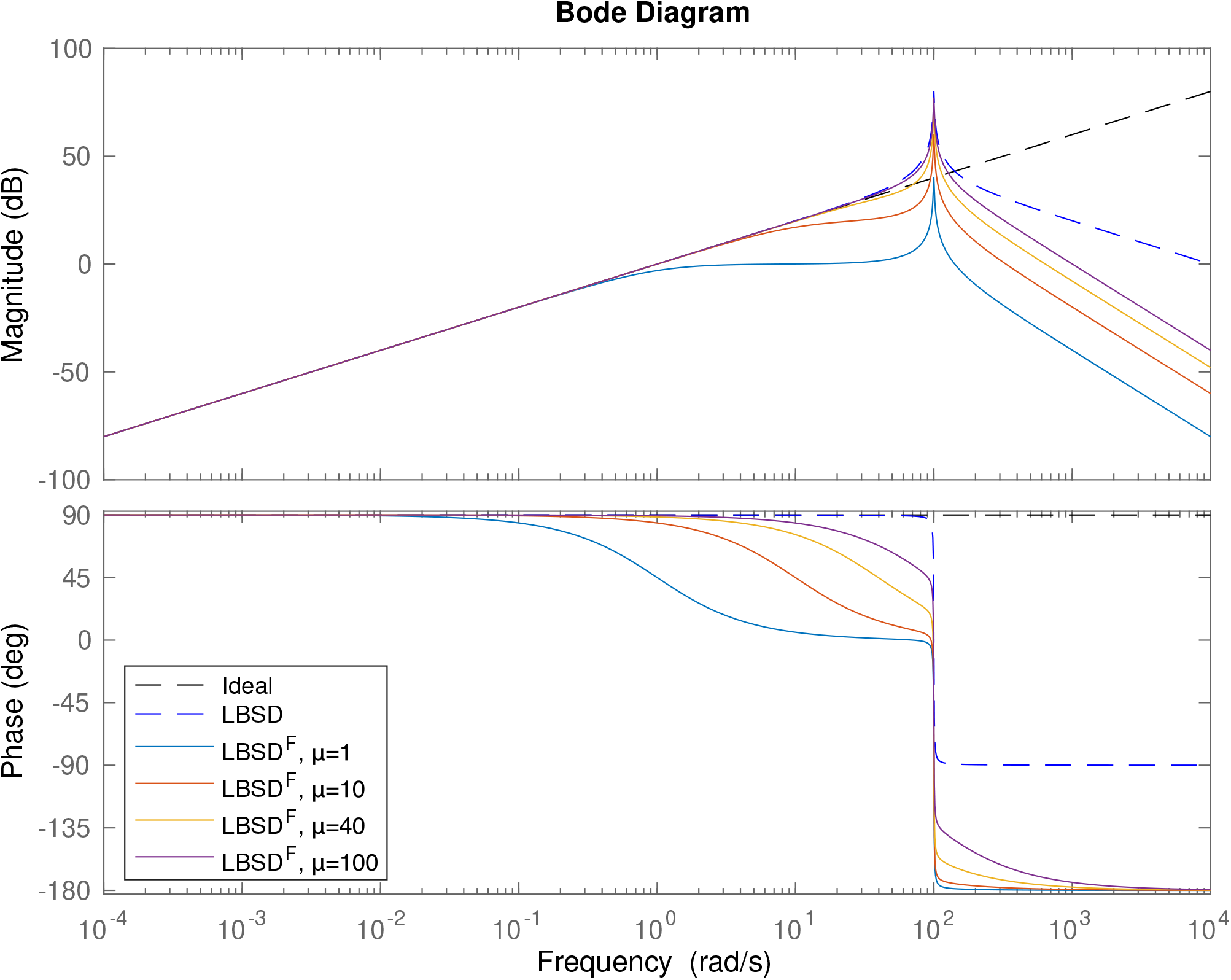
Frequency response analysis of Biomolecular Signal Differentiators^*F*^. Bode diagram depicting the magnitude and phase shift regarding the frequency response of a BioSD^*F*^ differentiator (Eq. (S83)) with *ε* = 10^−4^. We consider different values of *μ*, where *μ* = *μ*_1_ = *μ*_2_, that correspond to different colours. For comparison purposes, we also depict the bode plot (magnitude and phase) of the transfer function of an ideal differentiator (Eq. (S72)) and a BioSD differentiator (Eq. (S73)) with *ε* = 10^−4^ which are represented by black and blue dashed lines, respectively.

**Supplementary Figure 5:**
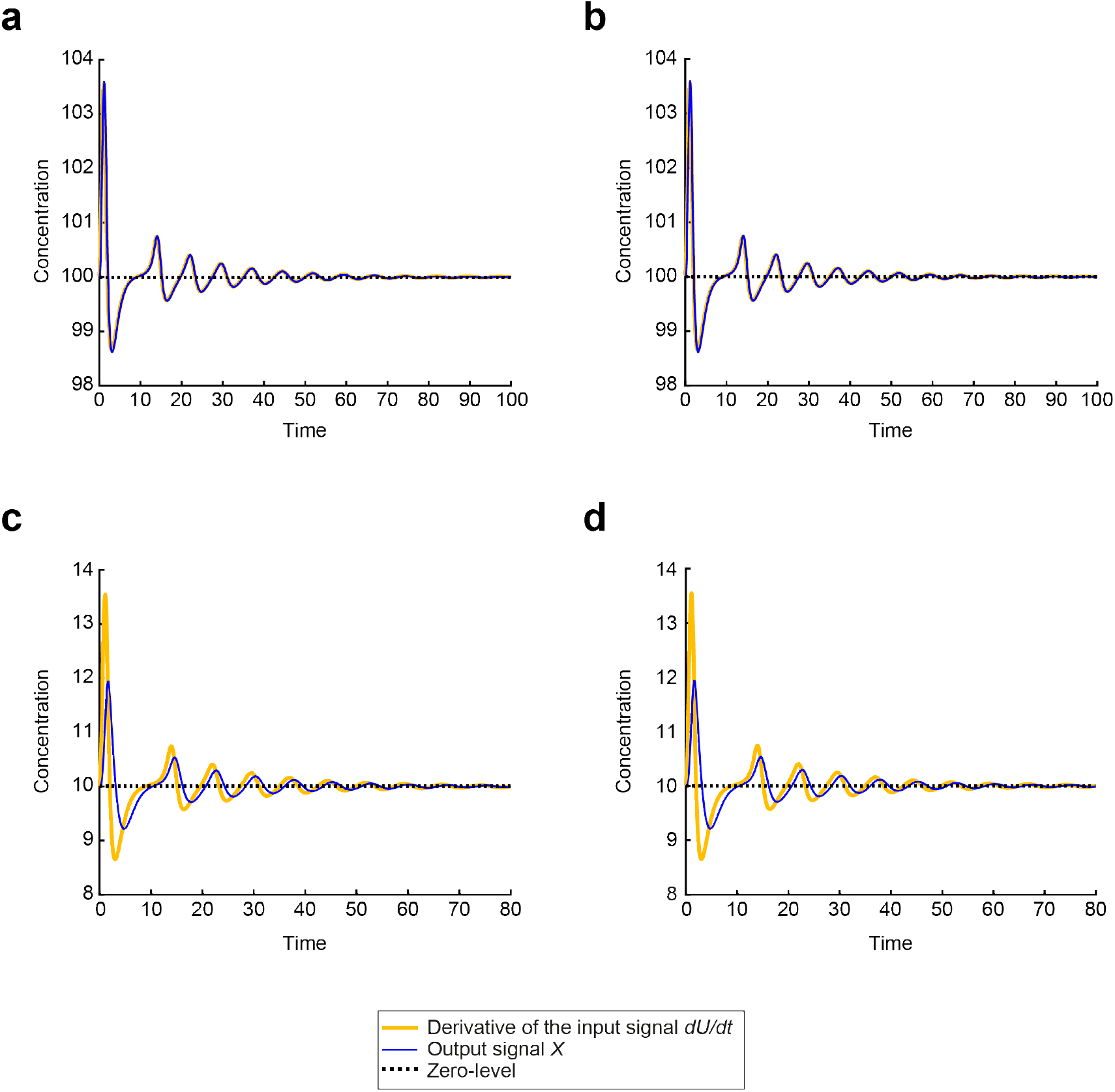
Sensing the rate-of-change of a synthetic regulatory biomolecular network through a Biomolecular Signal Differentiator. **a** Simulation of the BioSD-II (Eqs. (S11)–(S13)) response to the input presented in Fig. 2c with *η* = 3000, *b* = 150 and the remaining parameter values equal to those used in Fig. 2d. *η* can be characterized as sufficiently large since condition (S43) is satisfied. **b** Simulation of the BioSD-III (Eqs. (S23)–(S25)) response to the input presented in Fig. 2c with *η* = 30 and the remaining parameter values equal to those used in Fig. 2d. **c** The simulation in **a** is repeated with the values of *k_in_, k*_3_, *b* set to 10, 10 and 100, respectively (condition (S43) is satisfied). **d** The simulation in **b** is repeated with the values of both *k_in_* and *k*_3_ set to 10. In **a**, **b** condition (S60) is satisfied, i.e. *ε* ≪ 1. In contrast, *ε* ≫ 1 in **c**, **d**.

**Supplementary Figure 6:**
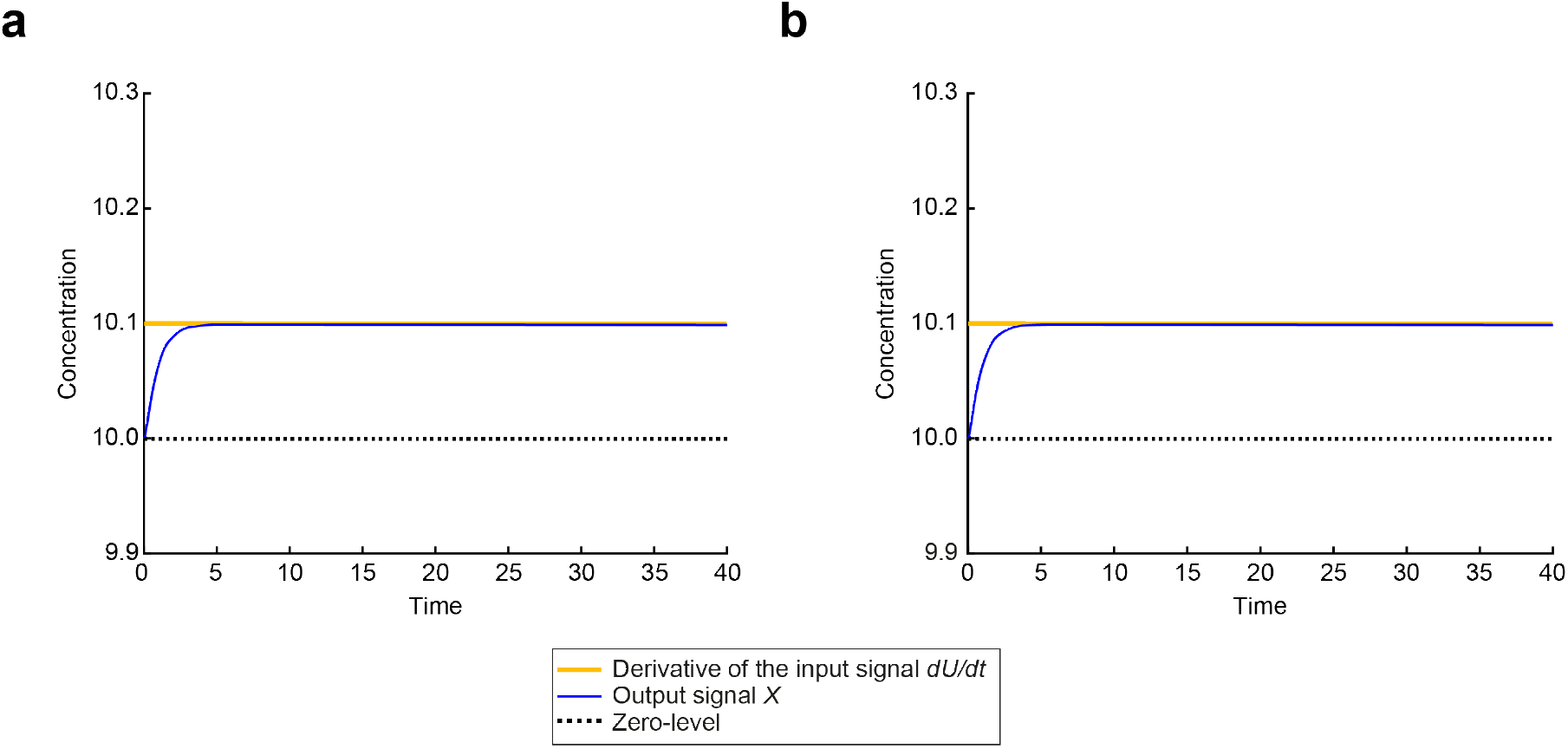
Sensing the rate-of-change of a birth-death biomolecular process through a Biomolecular Signal Differentiator. **a** Simulation of the BioSD-II (Eqs. (S11)–(S13)) response to the input presented in Fig. 3c with *η* = 3000 and the remaining parameter values equal to those used in Fig. 3d. *η* can be described as sufficiently large since condition (S43) is satisfied. **b** Simulation of the BioSD-III (Eqs. (S23)–(S25)) response to the input presented in Fig. 3c with *η* = 30 and the remaining parameter values equal to those used in Fig. 3d. In both **a** and **b**, condition (S60) is violated (*ε* ≫ 1.)

**Supplementary Figure 7:**
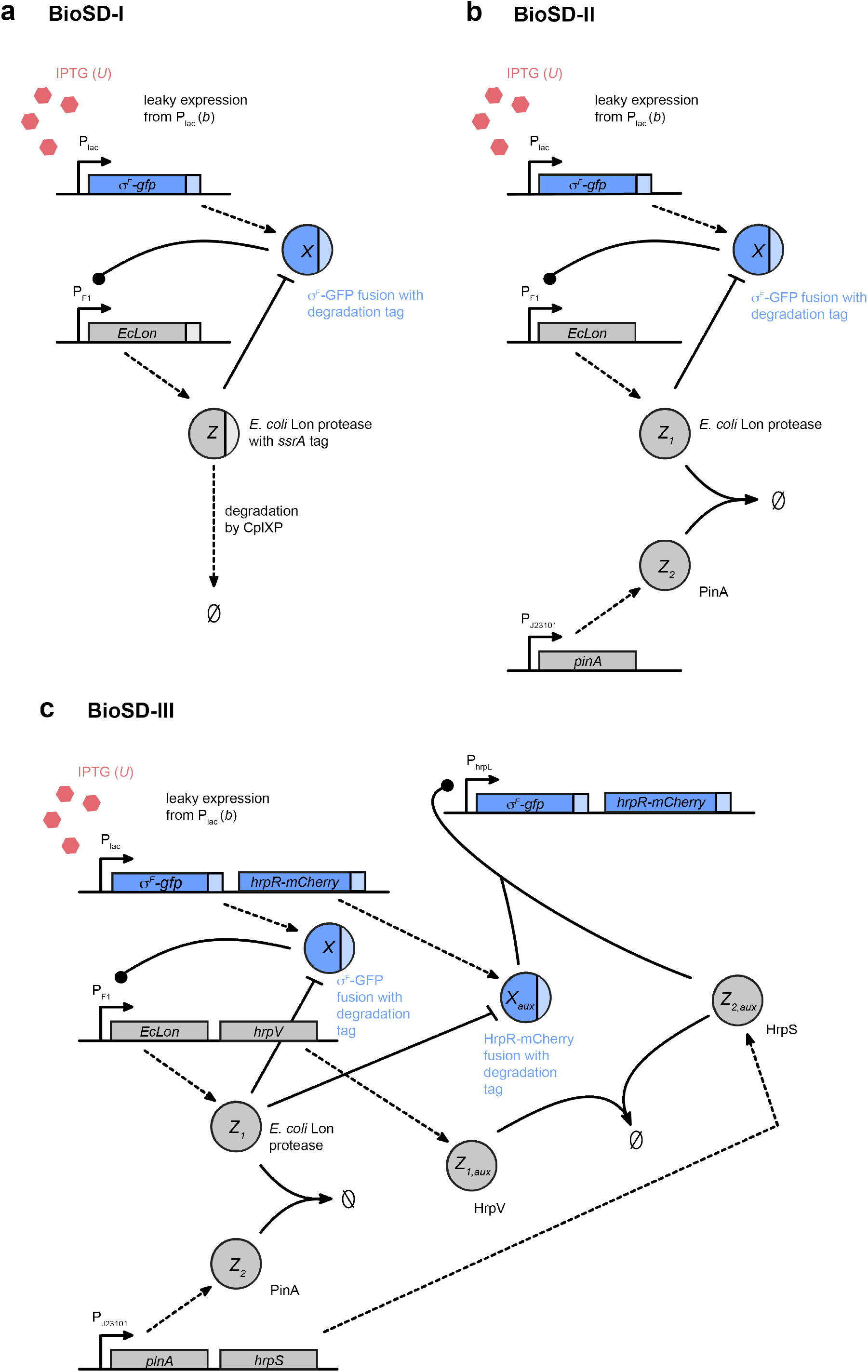
Possible experimental implementations of Biomolecular Signal Differentiators. Schematic representation of synthetic designs for **a** BioSD-I, **b** BioSD-II and **c** BioSD-III.

**Supplementary Figure 8:**
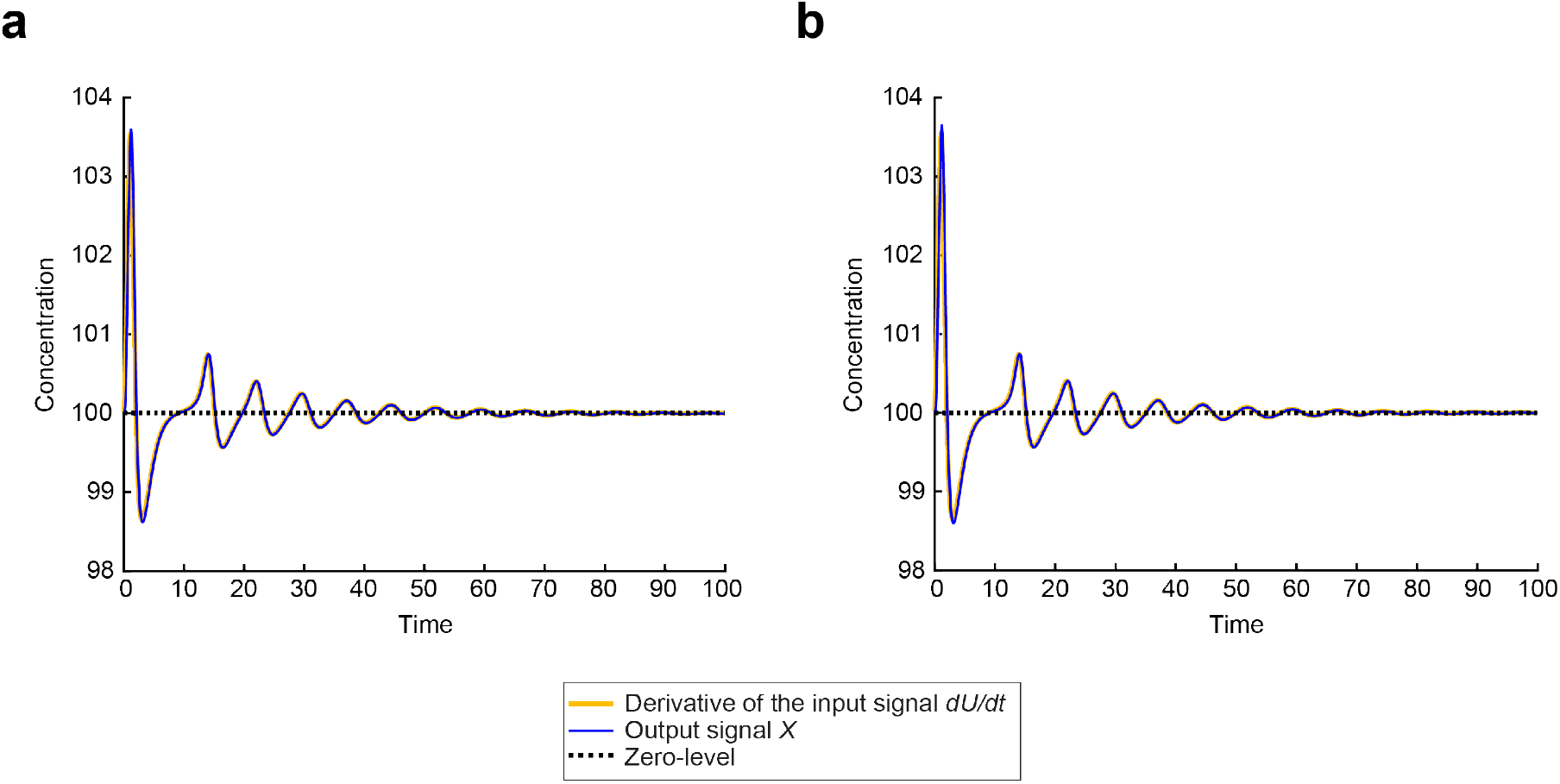
Sensing the rate-of-change of a synthetic regulatory biomolecular network through the proposed (experimental) circuit of Biomolecular Signal Differentiator-III. **a** Simulation of the circuit given by Eqs. (S85)–(S90) using the input presented in Fig. 3c and the following parameter values: *k_in_* = *k*_3_ = *b* = 100, *k*_1_ = *k*_1*a*_ = *k*_1*b*_ = *k*_2_ = 1, *η* = *η_a_* = 30, *δ* = *δ_a_* = 0.5 (this scenario corresponds to the simulation depicted in Supplementary Figure 5b). **b** We repeat the simulation in **a** with the values of *k*_1*a*_, *k*_1*b*_, *η_a_, δ_a_* set to 1.5 (increase by 50%), 1.25 (increase by 25%), 45 (increase by 50%), 0.75 (increase by 50%), respectively. It is evident that in both **a** (ideal case) and **b** the output, *X*, of the differentiator is an accurate replica of the derivative of input *U* - the loss of accuracy in **b** is negligible.

## Footnotes

1 Although we consider a sinusoidal *U^TV^*, this is without loss of generality as *U^TV^* can be thought of as a Fourier component of a more general signal.

## References

[1] Hugh R Wilson. Spikes, decisions, and actions: the dynamical foundations of neurosciences. Oxford University Press, 1999.

[2] Karl Johan Åström and Richard M Murray. Feedback systems: an introduction for scientists and engineers. Princeton University Press, 2021.

[3] Uri Alon. An introduction to systems biology: design principles of biological circuits. CRC press, 2019.

[4] Thomas S Shimizu, Yuhai Tu, and Howard C Berg. “A modular gradient-sensing network for chemotaxis in *Escherichia coli* revealed by responses to time-varying stimuli”. In: Molecular Systems Biology 6.1 (2010), p. 382.

[5] Pablo A Iglesias and Peter N Devreotes. “Navigating through models of chemotaxis”. In: Current Opinion in Cell Biology 20.1 (2008), pp. 35–40.

[6] Naama Barkai and Stan Leibler. “Robustness in simple biochemical networks”. In: Nature 387.6636 (1997), pp. 913–917.

[7] Steven M Block, Jeffery E Segall, and Howard C Berg. “Adaptation kinetics in bacterial chemotaxis”. In: Journal of Bacteriology 154.1 (1983), pp. 312–323.

[8] Robert M Macnab and Daniel E Koshland. “The gradient-sensing mechanism in bacterial chemotaxis”. In: Proceedings of the National Academy of Sciences 69.9 (1972), pp. 2509–2512.

[9] Mathieu Cloutier and Peter Wellstead. “The control systems structures of energy metabolism”. In: Journal of The Royal Society Interface 7.45 (2010), pp. 651–665.

[10] Edward J Hancock et al. “The interplay between feedback and buffering in cellular homeostasis”. In: Cell Systems 5.5 (2017), pp. 498–508.

[11] Harrison Steel et al. “Challenges at the interface of control engineering and synthetic biology”. In: 2017 IEEE 56th Annual Conference on Decision and Control (CDC). IEEE. 2017, pp.1014–1023.

[12] Domitilla Del Vecchio, Aaron J Dy, and Yili Qian. “Control theory meets synthetic biology”. In: Journal of The Royal Society Interface 13.120 (2016), p. 20160380.

[13] Timothy K Lu, Ahmad S Khalil, and James J Collins. “Next-generation synthetic gene networks”. In: Nature Biotechnology 27.12 (2009), p. 1139.

[14] Wolfgang Halter, Zoltan A Tuza, and Frank Allgöwer. “Signal differentiation with genetic networks”. In: IFAC-PapersOnLine 50.1 (2017), pp. 10938–10943.

[15] Wolfgang Halter, Richard M Murray, and Frank Allgöwer. “Analysis of primitive genetic interactions for the design of a genetic signal differentiator”. In: Synthetic Biology 4.1 (2019), ysz015.

[16] Christian Cuba Samaniego, Giulia Giordano, and Elisa Franco. “Practical differentiation using ultrasensitive molecular circuits”. In: 2019 18th European Control Conference (ECC). IEEE. 2019, pp. 692–697.

[17] Christian Cuba Samaniego, Jongmin Kim, and Elisa Franco. “Sequestration and delays enable the synthesis of a molecular derivative operator”. In: 2020 59th IEEE Conference on Decision and Control (CDC). IEEE. 2020, pp. 5106–5112.

[18] Michael Chevalier et al. “Design and analysis of a proportional-integral-derivative controller with biological molecules”. In: Cell Systems 9.4 (2019), pp. 338–353.

[19] Max Whitby et al. “PID control of biochemical reaction networks”. In: IEEE Transactions on Automatic Control (2021).

[20] Nuno MG Paulino et al. “PID and state feedback controllers using DNA strand displacement reactions”. In: IEEE Control Systems Letters 3.4 (2019), pp. 805–810.

[21] Kevin Oishi and Eric Klavins. “Biomolecular implementation of linear I/O systems”. In: IET Systems Biology 5.4 (2011), pp. 252–260.

[22] Saurabh Modi, Supravat Dey, and Abhyudai Singh. “Proportional and derivative controllers for buffering noisy gene expression”. In: 2019 IEEE 58th Conference on Decision and Control (CDC). IEEE. 2019, pp. 2832–2837.

[23] Maurice Filo and Mustafa Khammash. “A Class of Simple Biomolecular Antithetic Proportional-Integral-Derivative Controllers”. In: bioRxiv (2021).

[24] Domitilla Del Vecchio and Richard M Murray. Biomolecular feedback systems. Princeton University Press Princeton, NJ, 2015.

[25] Corentin Briat, Ankit Gupta, and Mustafa Khammash. “Antithetic integral feedback ensures robust perfect adaptation in noisy biomolecular networks”. In: Cell Systems 2.1 (2016), pp. 15–26.

[26] Corentin Briat, Ankit Gupta, and Mustafa Khammash. “Antithetic proportional-integral feedback for reduced variance and improved control performance of stochastic reaction networks”. In: Journal of The Royal Society Interface 15.143 (2018), p. 20180079.

[27] Noah Olsman, Fangzhou Xiao, and John C Doyle. “Architectural principles for characterizing the performance of antithetic integral feedback networks”. In: Iscience 14 (2019), pp. 277–291.

[28] Noah Olsman et al. “Hard limits and performance tradeoffs in a class of antithetic integral feedback networks”. In: Cell Systems 9.1 (2019), pp. 49–63.

[29] Noah Olsman and Fulvio Forni. “Antithetic integral feedback for the robust control of monostable and oscillatory biomolecular circuits”. In: IFAC-PapersOnLine 53.2 (2020), pp. 16826–16833.

[30] Ania-Ariadna Baetica, Yoke Peng Leong, and Richard M Murray. “Guidelines for designing the antithetic feedback motif”. In: Physical Biology (2020).

[31] Michael Samoilov, Adam Arkin, and John Ross. “Signal processing by simple chemical systems”. In: The Journal of Physical Chemistry A 106.43 (2002), pp. 10205–10221.

[32] Luca Laurenti et al. “Molecular filters for noise reduction”. In: Biophysical Journal 114.12 (2018), pp. 3000–3011.

[33] Aurelia Battesti, Nadim Majdalani, and Susan Gottesman. “The RpoS-mediated general stress response in *Escherichia coli*”. In: Annual Review of Microbiology 65 (2011), pp. 189–213.

[34] Regine Hengge-Aronis. “Signal transduction and regulatory mechanisms involved in control of the σ^*S*^ (RpoS) subunit of RNA polymerase”. In: Microbiology and Molecular Biology Reviews 66.3 (2002), pp. 373–395.

[35] Mihaela Pruteanu and Regine Hengge-Aronis. “The cellular level of the recognition factor RssB is rate-limiting for σ^*S*^ proteolysis: implications for RssB regulation and signal transduction in σ^*S*^ turnover in *Escherichia coli*”. In: Molecular Microbiology 45.6 (2002), pp. 1701–1713.

[36] Aurelia Battesti et al. “Anti-adaptors provide multiple modes for regulation of the RssB adaptor protein”. In: Genes & Development 27.24 (2013), pp. 2722–2735.

[37] David B Straus, William A Walter, and Carol A Gross. “The heat shock response of *E. coli* is regulated by changes in the concentration of σ^32^”. In: Nature 329.6137 (1987), pp. 348–351.

[38] Davide Roncarati and Vincenzo Scarlato. “Regulation of heat-shock genes in bacteria: from signal sensing to gene expression output”. In: FEMS Microbiology Reviews 41.4 (2017), pp. 549–574.

[39] David B Straus, William A Walter, and Carol A Gross. “DnaK, DnaJ, and GrpE heat shock proteins negatively regulate heat shock gene expression by controlling the synthesis and stability of σ^32^”. In: Genes & Development 4.12A (1989), pp. 2202–2209.

[40] Jürgen Gamer, Hermann Bujard, and Bernd Bukau. “Physical interaction between heat shock proteins DnaK, DnaJ, and GrpE and the bacterial heat shock transcription factor σ^32^”. In: Cell 69.5 (1992), pp. 833–842.

[41] David B Straus, William A Walter, and Carol A Gross. “The activity of σ^32^ is reduced under conditions of excess heat shock protein production in *Escherichia coli*”. In: Genes & Development 3.12A (1989), pp. 2003–2010.

[42] Pablo A Iglesias and Changji Shi. “Comparison of adaptation motifs: Temporal, stochastic and spatial responses”. In: IET Systems Biology 8.6 (2014), pp. 268–281.

[43] Elsa Bazellières et al. “Control of cell–cell forces and collective cell dynamics by the intercellular adhesome”. In: Nature Cell Biology 17.4 (2015), pp. 409–420.

[44] Stephanie K Aoki et al. “A universal biomolecular integral feedback controller for robust perfect adaptation”. In: Nature 570.7762 (2019), pp. 533–537.

[45] Hsin-Ho Huang, Yili Qian, and Domitilla Del Vecchio. “A quasi-integral controller for adaptation of genetic modules to variable ribosome demand”. In: Nature Communications 9.5415 (2018), pp. 1–12.

[46] Ciaran L Kelly et al. “Synthetic negative feedback circuits using engineered small RNAs”. In: Nucleic Acids Research 46.18 (2018), pp. 9875–9889.

## References

[1] Indra Bervoets et al. “A sigma factor toolbox for orthogonal gene expression in *Escherichia coli*”. In: Nucleic Acids Research 46.4 (2018), pp. 2133–2144.

[2] Wilson W Wong, Tony Y Tsai, and James C Liao. “Single-cell zeroth-order protein degradation enhances the robustness of synthetic oscillator”. In: Molecular Systems Biology 3.130 (2007), pp. 1–8.

[3] Jordan Ang et al. “Considerations for using integral feedback control to construct a perfectly adapting synthetic gene network”. In: Journal of Theoretical Biology 266.4 (2010), pp. 723–738.

[4] James J Hilliard, Michael R Maurizi, and Lee D Simon. “Isolation and characterization of the phage T4 PinA protein, an inhibitor of the ATP-dependent Lon protease of *Escherichia coli*”. In: Journal of Biological Chemistry 273.1 (1998), pp. 518–523.

[5] Jason R Kelly et al. “Measuring the activity of BioBrick promoters using an *in vivo* reference standard”. In: Journal of Biological Engineering 3.4 (2009), pp. 1754–1611.

[6] Stephanie K Aoki et al. “A universal biomolecular integral feedback controller for robust perfect adaptation”. In: Nature 570.7762 (2019), pp. 533–537.

[7] Baojun Wang, Mauricio Barahona, and Martin Buck. “Engineering modular and tunable genetic amplifiers for scaling transcriptional signals in cascaded gene networks.” In: Nucleic Acids Research 42.14 (2014), pp. 9484–9492.

